# Genome-wide association study reveals the genetic architecture of 27 yield-related traits in tomato

**DOI:** 10.1101/2020.10.01.322214

**Authors:** Jie Ye, Xin Wang, Wenqian Wang, Huiyang Yu, Guo Ai, Changxing Li, Pengya Sun, Xianyu Wang, Hanxia Li, Bo Ouyang, Junhong Zhang, Yuyang Zhang, James J. Giovannoni, Zhangjun Fei, Zhibiao Ye

## Abstract

Tomato (*Solanum lycopersicum*) is a highly valuable vegetable crop and yield is one of the most important traits. Uncovering the genetic architecture of yield-related traits in tomato is critical for the management of vegetative and reproductive development, thereby enhancing yield. Here we perform a comprehensive genome-wide association study for 27 yield-related traits in tomato. A total of 239 significant associations corresponding to 129 loci, harboring many reported and novel genes related to vegetative and reproductive development, were identified, and these loci explained an average of ∼8.8% of the phenotypic variance. A total of 51 loci associated with 25 traits have been under selection, especially during tomato improvement. Furthermore, a candidate gene, *SlALMT15* that encodes an aluminum-activated malate transporter, was functionally characterized and shown to act as a pivotal regulator of leaf stomata formation through increasing photosynthesis and drought resistance. This study provides valuable information for tomato genetic research and breeding.

---

Tomato (*Solanum lycopersicum*) is considered the leading vegetable crop, with a global production of 182.3 million tons in 2018 (FAOSTAT; http://www.fao.org/faostat), and has served as a model system for fleshy fruit biology. Tomato originated in South America^1^, but there is a long history of ‘wild’ tomato distribution in the mountainous area of southwest Guangxi of China. The natural barrier formed by the steep terrain ensures the independent evolution of these tomatoes, which contain valuable genetic resources of quality and resistance^2^.

Traits directly related to yield, such as fruit size, have been well studied^3, 4, 5, 6^. However, other yield-related traits, such as inflorescence location, inflorescence architecture, flower development and leaf stoma density, are also vital to the production in tomato. Most tomato varieties exhibit a sympodial growth habit instead of monopodial branching^7^, and domesticated tomatoes generate zigzag inflorescences with various flower numbers and inflorescence branches^8^. Taken together, flowering pattern, branching style and inflorescence polymorphism contribute to the total production in tomato.

Because of its importance, a number of genes controlling yield-related traits have been identified. *SP* (*SELF PRUNING*), homologous to *TFL1* (*TERMINAL FLOWER 1*) and *FT* (*FLOWERING LOCUS T*) from *Arabidopsis* and *CEN* (*CENTRORADIALIS*) from *Antirrhinum*, prevents flowering in the sympodial shoots and reduces the number of leaves only in the sympodial segments^7^. *LC* (*LOCULE NUMBER*), a WUSCHEL gene also named *lcn2*.*1*, has a major effect on the locule number of tomato fruit by controlling stem cell fate in the apical meristem^9^. *FAS* (*FASCINATED*), encoding a YABBY transcription factor, is considered a major gene and has the strongest effect in increasing the number of locules (from two to more than six)^10^. Moreover, *AN* (*ANANTHA*) and *S* (*COMPOUND INFLORESCENCE*) regulate inflorescence branches^11^, *SP5G* (*SELF PRUNING 5G*) promotes day-neutrality and early yield in tomato^12^, *SFT* (*SINGLE FLOWER TRUSS*) promotes flowering and attenuates apical meristem growth through systemic *SFT* signals^13^, *TMF* (*TERMINATING FLOWER*) has a key role in determining simple versus complex inflorescences^8^, *FA* (*FALSIFLORA*) causes highly branched inflorescences^14^, *Style 2*.*1* controls the style length in cultivated tomatoes^15^, *SlBOP1/2/3* (*BLADE-ON-PETIOLE*) promotes inflorescence complexity by interacting with *TMF*^16^, and *Jointless2* (*j2*) and *Enhancer-of-jointless2* (*EJ2*) are two homologs of the *Arabidopsis* floral organ identity MADS-box gene *SEPALLATA4* (*SEP4*) and regulate the formation of flower abscission zone^17^.

A goal of modern agriculture is to improve plant drought tolerance and production per amount of water used, referred to as water use efficiency (WUE)^18^. Stomata, epidermal structures that modulate CO_2_ and water vapor exchange between plants and the atmosphere, play critical roles in primary productivity and in plant adaptation to the global climate. Positively acting transcription factors and negatively acting mitogen-activated protein kinase (MAPK) signaling control stomatal development in Arabidopsis^19, 20, 21, 22^; however, the key regulatory genes of stomata formation in tomato haven’t been cloned yet.

Genome-wide association study (GWAS) is an effective approach to investigate the genetic architecture of complex agronomic traits in crops^23, 24, 25, 26, 27^. In tomato, using bi-parental populations, several loci for important agronomic traits have been identified^28^. Recently, GWAS has been employed to explore genetic loci associated with 15 agronomic traits in 163 tomato accessions^29^. Here, we improve the tomato haplotype map by adding sequencing data of 66 Guangxi tomatoes and present a GWAS for 27 agronomic traits to identify a substantial number of loci potentially associated with tomato production. Several genomic loci underlying these agronomic traits are consistent with previous reports and many loci are newly identified in this study. We further functionally verified a candidate gene underlying the GWAS signal of leaf stomatal density.

## RESULTS

### Sequencing, variants and population structure of Guangxi tomatoes

In this study, a total of 605 tomato accessions were used for genotyping and subsequent GWAS analysis. Among these accessions, a diverse global collection of 539 accessions were genotyped in previous studies ^30, 31, 32^, and 66 newly sequenced accessions, which showed vines and had small leaves and small red fruits with thin skin, were collected from the mountainous area of Guangxi Province, China (**Supplementary Table 1** and **Supplementary Fig. 1**). A total of 6.25 billion 100-bp paired-end reads were obtained for these Guangxi (GX) accessions, representing a mean depth of 9.85× coverage of the tomato genome (**Supplementary Table 1**). A total of 4,412,112 million high-quality SNPs were called from the sequencing data of the 605 accessions.

**Table 1.** Summary of significant locus-trait associations identified in GWAS.

| Item | Organ location<br>traits | Organ number<br>traits | Organ size<br>traits |
| --- | --- | --- | --- |
| Number of traits | 4 | 9 | 14 |
| Number of suggestive SNPs (significant<br>SNPs)<sup>a</sup> | 28 (18) | 85 (49) | 126 (81) |
| Number of loci per trait | 4-11 | 2-18 | 1-20 |
| Lead SNPs explaining >15% of variation<sup>b</sup> | 7 | 10 | 14 |
| Maximum explained variation (%) | 69.4 | 78.5 | 75.3 |
| Explained variation per SNP (%) | 9.8 | 8.0 | 7.9 |

<sup>a</sup>Suggestive SNPs, $P \leq 2.4 \times 10^{-7}$ ; Significant SNPs, $P \leq 1.2 \times 10^{-8}$ .
<sup>b</sup>Lead SNPs are those with the lowest $P$ values in the defined association loci.

**Figure 1.**
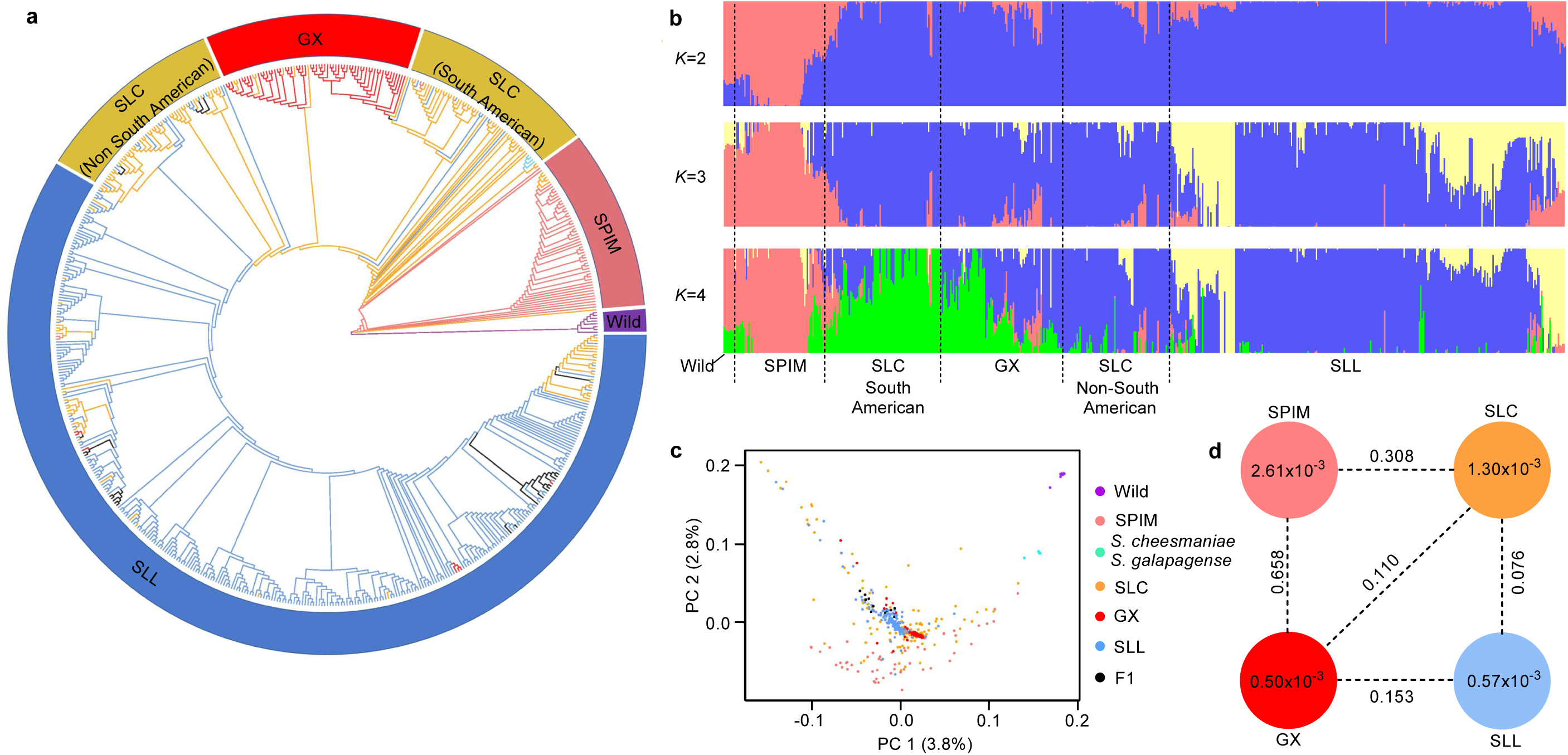
Population diversity of the tomato accessions. (**a**) Neighbor-joining phylogenetic tree of 605 tomato accessions. The outer ring indicates different groups. Colors of branches on the tree indicate different groups: SPIM (pink), SLC (orange), GX (red), SLL (blue), and other wild (purple). (**b**) Population structure of tomato accessions with different numbers of clusters (*K* = 2, 3 and 4). The orders and positions of the accessions on the *x* axis are consistent with those in the neighbor-joining tree. (**c**) PCA plot of the tomato accessions. The dot color scheme is same as indicated in **a**. PC1, first principal component; PC2, second principal component. (**d**) Nucleotide diversity (π) and population divergence (*F*_ST_) among the four groups. For each group, nucleotide diversity (π) is shown inside the circle. Population divergence (*F*_ST_) between the two groups is shown on the dotted line.

To determine the evolutionary status of the GX tomatoes, we inferred the phylogenetic relationships of the 605 tomato accessions using the genome-wide SNPs. Largely consistent with the previously reported phylogeny^30^, these tomatoes were divided into six groups (**Fig. 1a**), which was further supported by the principal component analysis (PCA) and population structure analysis (**Fig. 1b,c**). Interestingly, GX tomato accessions clustered together and divided the *S. lycopersicum* var. *cerasiforme* (SLC) group into two subgroups: South American SLC and Non-South American SLC, suggesting that the GX accessions were introduced to China from South America after domestication and then underwent independent evolution.

The nucleotide diversity decreased from the *S. pimpinellifolium* (SPIM) group (π=2.61×10^−3^) to the SLC group (π=1.3×10^−3^) and to the *S. lycopersicum* var. *lycopersicum* (SLL) group (π=0.57×10^−3^) and GX group (π=0.5×10^−3^), indicating that a large amount of genetic diversity has been lost for GX tomatoes, possibly because of geographical isolation and narrow ancestral genetic background. Large population divergences between the GX group and the other three groups were observed (**Fig. 1d**), suggesting that GX tomatoes in China might have accumulated some genetic diversity after their introduction.

### Phenotypic variation in the tomato population

In this study, a total of 27 yield-related traits, including fruit number in the second spike (FRNS), fruit number in the third spike (FRNT), flower number on the second inflorescence (FLNS), flower number on the third inflorescence (FLNT), sepal number (SN), petal number (PN), first inflorescence node (FIN), ovary transverse diameter (OTD), ovary longitudinal diameter (OLD), ovary transverse diameter to ovary longitudinal diameter ratio (OTLD), stigma exsertion (SE), stigma shape (SS), sepal length to petal length ratio (SPR), stomatal density (SD), first to second inflorescence node (FSIN), internode length (IL), petal length (PL), stamen length (STAL), stigma length (STIL), sepal length (SL), indeterminate or determinate meristem (IDM), stamen length to (stigma length+ovary longitudinal diameter) ratio (SSR), fruit stalk diameter (FSD), fruit stalk length (FSL), inflorescence type (IT) and fasciated flower (FL), were investigated during the whole growth period of tomato with three replications (**Supplementary Fig. 2** and **Supplementary Note**). These agronomic traits were classified into three categories: four organ location traits, nine organ number traits and 14 organ size traits (**Supplementary Tables 2** and **3**). For the majority of the agronomic traits, an abundant variation was detected, with the coefficients of variation (*CV*) ranging from 0.12 for PN to 2.92 for FLNS (**Supplementary Table 2**). All traits determined in the diverse global collection of tomato accessions (**Supplementary Table 3**) displayed a broad-sense heritability (*H*^*2*^) greater than 0.7, and 14 had heritability over 0.9 (**Supplementary Table 2**), suggesting these traits were primarily determined by genotype. Most of the traits showed a normal distribution while PN, SN, FSIN, FSD, FSL and OTD showed skewed distributions (**Supplementary Fig. 3**). Except IL, the phenotypic values of all agronomic traits were significantly different among three subgroups of tomato (SPIM, SLC and SLL) (**Supplementary Fig. 4**). For example, the ovary size (OLD and OTD) increased during tomato domestication and improvement, while the flower number (FLNS and FLNT) and fruit number (FRNS and FRNT) declined (**Supplementary Fig. 4**). Many of the 27 traits were correlated (**Supplementary Fig. 5**), and the correlation coefficients between some traits were very high, such as SL and PL (*r* = 0.85), IL and FSIN (*r* = 0.88), and OTD and SN (*r* = 0.83). Negative correlation between ovary size (OLD and OTD) and flower and fruit number (FLNS, FLNT, FRNS and FRNT) supported the hypothesis that fruit size was positively selected, and the number of fruits decreases during the process of tomato breeding (**Supplementary Fig. 5**).

### Genome-wide association study for 27 yield-related traits

To reveal the genetic architecture of the vegetative and reproductive organ development in tomato, we performed GWAS on the 27 yield-related traits in the 605 tomato accessions using the genome-wide SNPs. The Manhattan plots of GWAS for all 27 traits are shown in **Supplementary Figures 6-32**, and detailed information about all significant associations is summarized in **Supplementary Table 4**.

In tomato, an association locus has been defined as a chromosomal region in which the distance between the adjacent pairs of associated SNPs is less than 200 kb^33^. According to this definition, a total of 239 suggestive associations (including 148 significant associations) corresponding to 129 loci were identified (**Fig. 2** and **Supplementary Table 4**). For organ location, organ number and organ size traits, 28, 85, and 126 associated loci were identified, respectively (**Table 1**). On average, each trait had ∼9 identified associated loci, with SSR having 20 associated loci whereas SPR only one. We identified four potential GWAS hotspots (density>0.03) which fit perfectly with known QTLs involved in the regulation of the growth and development of tomatoes on chromosomes 2 (*LC*, locule number), 3 (*FA*, falsiflora), 6 (*SP*, self-pruning), and 11 (*FAS*, fascinated), respectively (**Supplementary Fig. 33**). For example, the locus overlapping with *LC* was simultaneously detected for 11 different traits including FLNS, FRNS, FLNT, FRNT, PN, SN, FSD, OTD, OTLD, SS and SSR. The locus overlapping with *FAS* was detected for 8 different traits including FSIN, IL, IDM, IT, PN, SN, FL, OTD and SS. These results are in line with that many of the agronomic traits were highly correlated (**Supplementary Fig. 5**). The percentage of phenotypic variation explained by each locus ranged from 1.6 to 69.4% in organ location traits, from 1.2 to 78.5% in organ number traits and from 1.4 to 75.3% in organ size traits, with mean values of 9.8%, 8.0% and 8.6%, respectively (**Table 1**). Although some traits were controlled by one major locus that explained over 60% of the natural variation, such as SN, IDM, IT and FL, most agronomic traits were determined by multiple moderate-effect loci.

**Figure 2.**
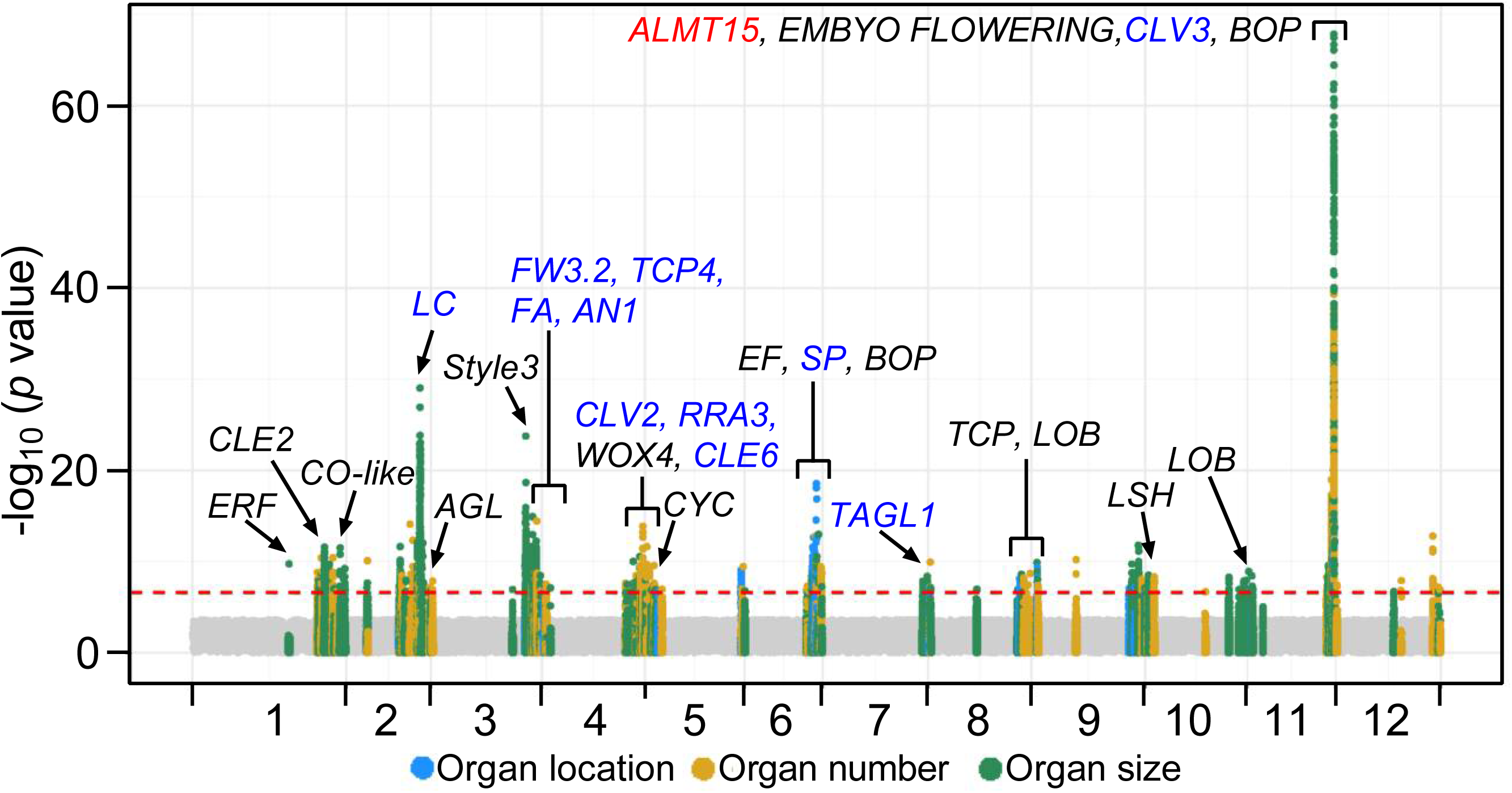
Manhattan plot of GWAS with identified genetic associations. The significance of the associations with agronomic traits is indicated as the negative logarithm of the *P* values. All trait-SNP associations with *P* values < 2.4×10^−7^ are plotted against the genome at 1-Mb intervals. Red horizontal dashed line indicates the genome-wide significance threshold (*P* values < 1.2×10^−8^). Associations with organ location traits are indicated by light blue circles, organ number traits by yellow circles, and organ size traits by green circles. Candidate genes including those with known functions (blue text) and unknown functions (black text) are indicated. Function of *ALMT15* in the red text was validated in this study.

### Key candidate genes involved in vegetative and reproductive development

We searched for functional known or unknown candidate genes responsible for the variation of tomato agronomic traits based on the information of gene annotation, phylogenetic analysis of candidate genes with their homologs with known functions, and cross-referencing with results from previous linkage mapping. We were able to identify several plausible candidate genes and possible causative SNPs underlying the yield-related traits (**Supplementary Table 5**). Taking the 18 associated loci of PN as an example, in addition to previously reported large-effect genes including *CLV3, LC, SlWOX4* and *SlRRA3*^34, 35^, we also identified several new minor-effect candidate genes such as *SlCLE2* and *SlCLE6* belonging to the CLV3/EMBRYO-SURROUNDING REGION family that plays an important role in regulating stem cell proliferation and differentiation of plant development^36^ (**Supplementary Fig. 34**). The new loci identified here provide valuable candidates for future studies that can further our understanding of the genetic regulation of these traits.

For FSIN, IL, PL, STAL and IDM, the association signal at the end of chromosome 6 showed a subtle zigzag pattern, suggesting multiple trait-associated genes present in this small region (**Fig. 3a**). In detail, we detected three significantly associated SNPs (SL2.50ch06_43765964, SL2.50ch06_44230173 and SL2.50ch06_45972263) corresponding to genes *Solyc06g071140, Solyc06g071830* and *Solyc06g074350* in this region, respectively (**Fig. 3b**). *Solyc06g074350* corresponds to the known *SELF-PRUNING* (*SP*) gene^7^. *Solyc06g071140* (*SlELF3*), located 28.7 kb downstream of SL2.50ch06_43765964, was annotated as a homolog of the EARLY FLOWERING gene that regulates circadian clock function and flowering in Arabidopsis^37, 38^, rice^39^ and pea^40^ (**Supplementary Fig. 35a**). *Solyc06g071830* (*SlBOP4*), located 15.3 kb downstream of SL2.50ch06_44230173 and encoding a BTB/POZ protein, was homologous to three other SlBOPs (*SlBOP1, SlBOP2* and *SlBOP3*) that are known to control inflorescence architecture and flower production in tomato^16^ (**Supplementary Fig. 35b**), suggested that *Solyc06g074350* is likely the candidate gene underlying this locus. To compare association results across different traits, we constructed an association network to visualize complex trait relationships (**Fig. 3c**). The SNP SL2.50ch06_45972263 near the *SP* gene was the most significant locus for the IDM trait (*P*=3.36×10^−55^) and was repeatedly detected for IL (*P*=3.05×10^−16^) and FSIN (*P*=8.63×10^−19^). The SNP SL2.50ch06_44230173 near *Solyc06g071830* was the second most significant locus of FSIN (*P*=2.28×10^−13^) and was repeatedly detected for IL (*P*=2.85×10^−11^), STAL (*P*=2.02×10^−10^) and PL (*P*=4.26×10^−10^). The SNP SL2.50ch06_43765964 near *Solyc06g071140* was the third most significant locus for FSIN (*P*=2.74×10^−12^) and was repeatedly detected for IL (*P*=2.72×10^−11^), STAL (*P*=1.17×10^−10^) and PL (*P*=8.5×10^−9^). The co-localization between these two organ size traits (PL and STAL) and three organ location traits (IL, FSIN and IDM) suggests a relationship between flowering time and flower size in tomato.

**Figure 3.**
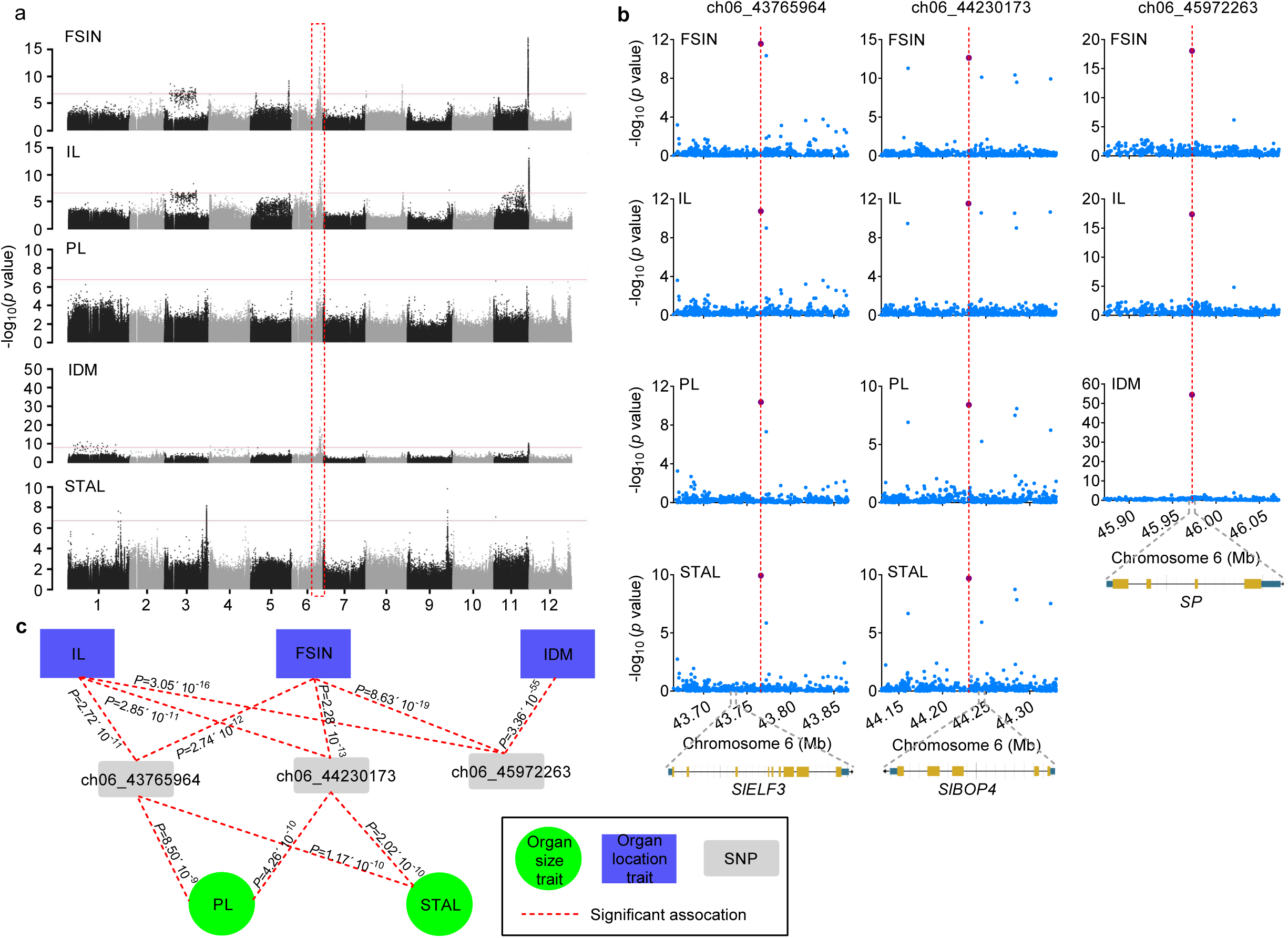
Genome-wide associations for FSIN, IL, PL, IDM and STAL. (**a**) Manhattan plots of GWAS for FSIN (first to second inflorescence node), IL (internode length), PL (petal length), IDM (indeterminate or determinate meristem) and STAL (stamen length). The common association signal is highlighted with a dashed red box. (**b**) Regional (200-kb) association plots of GWAS for FSIN, IL, PL, STAL and IDM. Three common significant association SNPs (SL2.50ch06_43765964, SL2.50ch06_44230173 and SL2.50ch06_45972263) are highlighted with dashed lines. Corresponding candidate genes *SlELF3* (*Solyc06g071140*), *SlBOP4* (*Solyc06g071830*) and *SP* (*Solyc06g074350*) are indicated. (**c**) Association network summarizing GWAS results shown in (**b**). GWAS results of certain trait containing any of the three significant SNPs are treated as nodes and are connected. Green circle indicates organ size traits (PL and STAL) and purple rectangle indicates organ location traits (IL, FSIN and IDM).

Stigma exsertion, defined as the pistil longer than the stamen (**Supplementary Note**), has been reported as a key determinant of the plant mating system and contributes to the application of heterosis in plant^41, 42^. *Style2*.*1*, the major QTL responsible for SE in cultivated tomatoes, has been identified on chromosome 2 (Ref. ^15^). In this study, SE and SSR were significantly associated with SL2.50ch01_84029382 (*P*=2.78×10^−12^) and SL2.50ch03_60427735 (*P*=1.29×10^−16^) on chromosome 1 and 3, respectively (**Supplementary Figs. 26** and **30**). SL2.50ch03_60427735 is located 2 bp from the start codon of *Solyc03g098070* that encodes an C2H2L domain class transcription factor (referred to as *Style3* hereafter). Homologous of *Style3* in *Arabidopsis*, SGR5, has been reported to regulate the gravitropism of inflorescence stems^43^. Both association and expression analyses supported that *Style3* is likely the candidate gene underlying stigma exsertion in tomato (**Supplementary Fig. 36**). *Style3* was highly expressed in flower tissues, especially in style (**Supplementary Fig. 36c**). Higher expression of *Style3* was found in stigma exsertion and stigma flush tomatoes compared with stigma inside tomatoes, while no significant expression difference was observed between stigma exsertion and stigma flush accessions (**Supplementary Fig. 36d**). Genotyping analysis revealed that all stigma exsertion and stigma flush accessions (52 SLC, 22 SLL, 18 SPIM and 11 other wild accessions) exhibited the C allele, while the stigma inside accessions (6 SLC and 37 SLL accessions) exhibited the T allele of the lead SNP (SL2.50ch03_60427735; **Supplementary Fig. 36e** and **Supplementary Table. 6**). These results provide the first evidence that *Style3* may be involved in determining the style length.

Despite the widely reported genetic mapping of the fruit size trait, genes responsible for early fruit development remain largely unexplored in tomato^5, 6, 44^. For the GWAS of ovary size traits, four clear signals, SL2.50ch01_84023965 corresponding to *fw1*.*2*, SL2.50ch02_47188498 corresponding to *lc*, SL2.50ch03_64734105 corresponding to *fw3*.*2* and SL2.50ch11_55052389 corresponding to *fas*, were identified for OTD and OTLD, but not for OLD, indicating that lateral development in early fruit stages is more genetically regulated in tomato (**Supplementary Figs. 23** and **24**).

### Functional characterization of a candidate gene regulating stomatal density

Stomata, the exchange channels of gas and moisture between the plant and the external atmosphere, is an important trait affecting photosynthesis, transpiration and productivity of plants^19^. Here, we experimentally verified and further characterized a candidate gene involved in stomatal formation, providing novel functional insights. The phenotype values of stomatal density presented a skewed normal distribution in the natural tomato population investigated here (**Supplementary Fig. 3**). We obtained two significant loci associated with stomatal density in tomato leaf on chromosome 3 and 11, respectively (**Supplementary Fig. 17**). The significant association (*P*=5.59×10^−8^) between SNP SL2.50ch11_53544569 and stomatal density suggested that a genomic sequence related to SNP SL2.50ch11_53544569 forms the major genetic locus (explain 26.2% of the variation) responsible for the natural variation in stomatal formation of tomato leaves (**Fig. 4a**). Two major haplotypes, C and T, at the lead SNP (SL2.50ch11_53544569) of the association signal were associated with high-density and low-density stomatal phenotypes in tomato, respectively (**Fig. 4b**).

**Figure 4.**
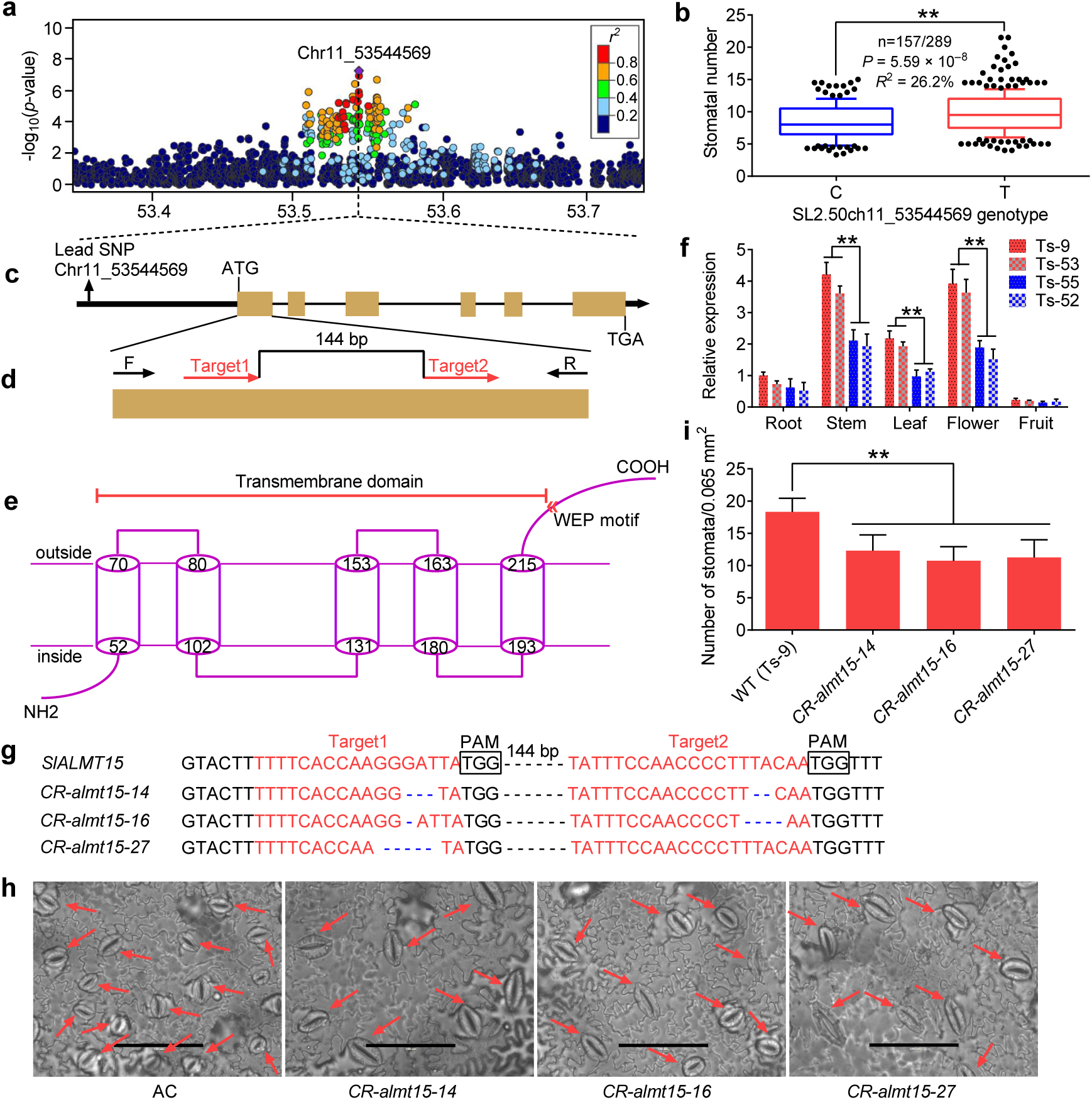
Genetic associations with stomatal density and functional characterization of *SlALMT15*. (**a**) Locus on chromosome 11 associated with stomatal density identified through GWAS. Regional Manhattan plot of a genomic region spanning 400 kb centered at the peak SNP is shown. Lead SNP is indicated in purple. A representation of pairwise *r*^*2*^ values (a measure of LD) between the lead SNP and all other SNPs in this 400-kb region is shown, where the color of each dot corresponds to the *r*^*2*^ value according to the color scale. (**b**) Box plot of stomatal densities in tomato accessions with different alleles (C or T) at SNP ch11_53544569. (**c**) Gene structure of *SlALMT15*. Filled black, filled orange and black lines represent promoter, coding sequence and introns, respectively. (**d**) Schematic illustration of the two sgRNA target sites (red arrows) in *SlALMT15*. Black arrows represent the location of PCR genotyping primers. (**e**) Proposed topology for SlALMT15. The N-terminal contains five transmembrane domains (cylinder). The position of the highly conserved WEP-motif at the C-terminal is indicated. (**f**) Transcript levels of *SlALMT15* in different tomato tissues (root, stem, leaf, flower and red fruit). Ts-9 and Ts-53 are high-density stomata accessions, whereas Ts-52 and Ts-55 are low-density stomata accessions. (**g**) Mutated alleles identified from three T_2_ *CR-almt15* mutant lines. Red letters indicate sgRNA target sequences, and black boxes indicate protospacer-adjacent motif (PAM) sequences. (**h**) Images showing the abaxial epidermis of leaves from wild type and three *CR-almt15* mutant lines. Red arrows point to stomata. Scale bar, 100 μm. (**i**) Number of stomata per unit area in leaves of wild type and three *CR-almt15* lines. Stomatal number was counted in each field of view (200×, ∼0.065 mm^2^) of three plants. The third leaf from the top of 6-weeks-old transgenic and non-transgenic plants was used for stomata number analysis. Asterisks indicate significant differences by *t* test: \*\**P*<0.01.

There were 22 genes within the 100-kb sequences flanking either side of the lead SNP (**Supplementary Table 7**). Pairwise linkage disequilibrium (LD) analysis within the 400-kb interval centered on the lead SNP showed that SNPs with high LD to the lead SNP fell into a 70-kb region for 53.51 Mb to 53.58 Mb (**Fig. 4a**). *Solyc11g068970*, encoding an aluminum-activated malate transporter (ALMT) (**Fig. 4c**), was the closest gene to the lead SNP (1.8 kb downstream), and the gene and the SNP were in the same LD block (**Supplementary Fig. 37**). Previous studies have shown that many ALMTs are expressed in guard cells and contribute to stomatal closure in plants^45, 46^, therefore, *Solyc11g068970* (which we named *SlALMT15*) was considered as the causal gene candidate for controlling stomatal density in tomato.

A total of 98 orthologs with high amino acid similarity (>50%) to SlALMT15 were identified in different plant species (**Supplementary Fig. 38**). The SlALMT15 protein was predicted to contain five transmembrane helices and a long C-terminal domain that harbored a conserved WEP-motif (**Fig. 4e**). To investigate functional allelic variation at the *SlALMT15* locus, we analyzed the nucleotide sequence of *SlALMT15* in 13 tomato accessions with diverse stomatal density, which revealed 17 polymorphisms including five indels and 12 SNPs in the promoter region, and no polymorphism in the gene region (**Supplementary Fig. 39**). Except two indels, the remaining 15 polymorphisms led to 36 possible *cis* element changes in the promoter of *SlALMT15* according to PLACE (https://www.dna.affrc.go.jp/PLACE/) (**Supplementary Table 8**). The spatial and temporal expression patterns of *SlALMT15* in high-density stomata accessions (Ts-9 and Ts-53) and low-density stomata accessions (Ts-55 and Ts-52) were then investigated. *SlALMT15* showed high expression levels in stem, flower and leaf, but low in fruit and root, with the transcript levels higher in most tissues of Ts-9 and Ts-53 than in Ts-55 and Ts-52 (**Fig. 4f**), supporting a role of *SlALMT15* in positively regulating stomatal density in tomato.

To further functionally characterize the role of *SlALMT15* and stomatal formation, we mutated *SlALMT15* in vivo using CRISPR/Cas9 in the high-density stomatal accession Ts-9 (**Fig. 4d**). CRISPR/Cas9-induced knockout mutations (deletions) in *SlALMT15* were detected by PCR and further confirmed by DNA sequencing (**Fig. 4g**). The three investigated mutant lines developed significantly less stomata than Ts-9 in leaves (**Fig. 4h** and **4i**). To investigate whether the *SlALMT15* affects drought stress tolerance by affecting stomatal density, six-week-old seedlings from the four studied mutant lines and the wild-type were challenged with drought stress by withholding water for 8 days. Dehydration symptoms (leaf wilting) were observed in both mutants and wild-type plants, but the wilting was significantly more severe in the wild-type plants. Changes of drought-related physiological indicators, including net photosynthetic rate, stomatal conductance, transpiration rate and malondialdehyde (MDA) content, also supported the different degrees of wilting in the mutants and the wild-type plants (**Supplementary Fig. 40**). Together these results strongly support that SlALMT15 functions in the stomata formation and further affects drought stress tolerance in tomato.

### Selective sweeps related to yield-related traits

Long-term domestication and improvement have brought many morphological changes to tomato, such as larger flower and fruit^6^, stronger stem^47^, embedded stigma^15^ and so on. To investigate how artificial selection underlying these changes, we scanned genomic regions for selective signals in the tomato genome. Based on the phylogenetic analysis, we combined the GX group and the SLC group into one group (SLC_GX) for selective sweep identification. In total, we identified 128 domestication sweeps exhibiting lower nucleotide diversity in SLC_GX compared with SPIM, covering 59.85 Mb and harboring 2,492 genes (π_SPIM_/π_SLC_GX_> 2.76; **Fig. 5a, Supplementary Fig. 41** and **Supplementary Table 9**). On the other hand, 204 improvement sweeps with a cumulative size of 68.22 Mb were detected and harbored 4,959 genes (π_SLC_GX_/π_SLL_> 5.38; **Fig. 5b** and **Supplementary Table 10**). Collectively, there were 2,132 and 4,599 genes only involved in the domestication or improvement, respectively, and 360 genes in both (**Supplementary Table 11**). We found that 62% of genes in domestication sweeps (1,545 out of 2,492) and 63% in improvement sweeps (3,122 out of 4,959) detected in our study were also detected in a previous study^30^ (**Fig. 5c**).

**Figure 5.**
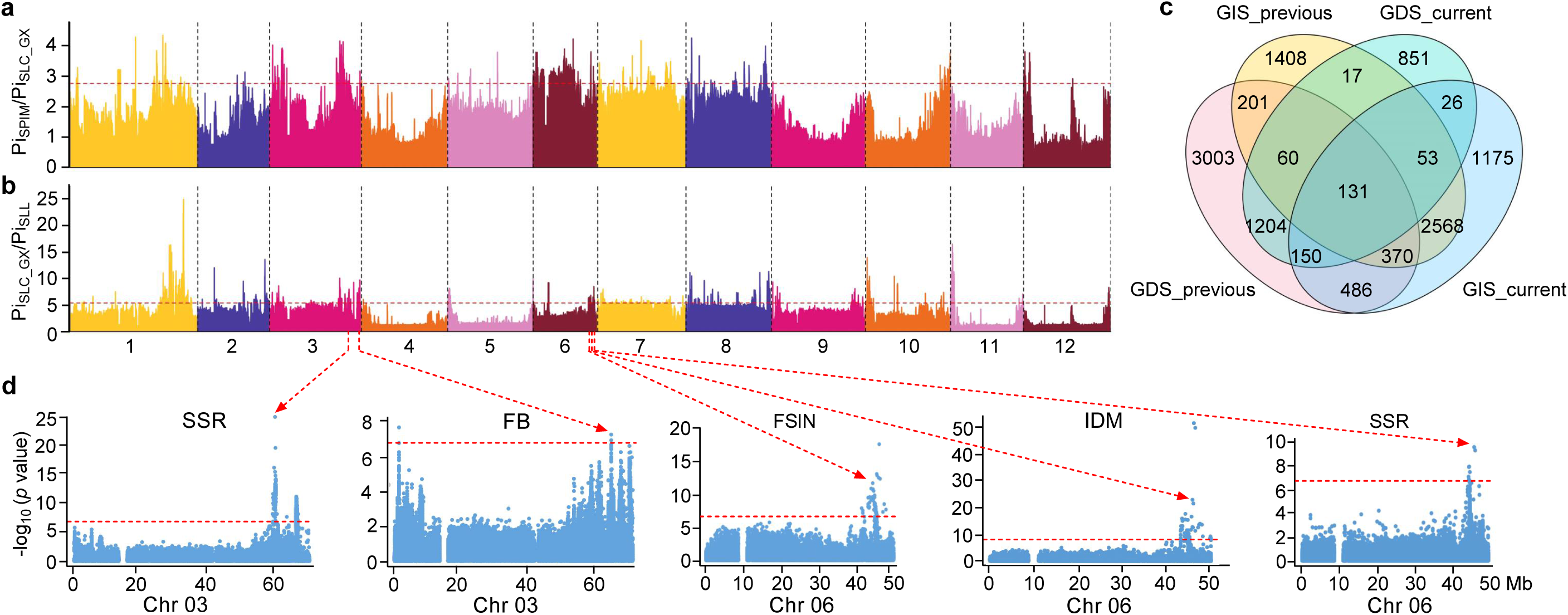
Genome-wide screen of selective sweeps during tomato domestication and improvement. (**a, b**) Selection signals during tomato domestication (**a**) and improvement (**b**). The horizontal red dashed lines indicate the genome-wide threshold for domestication sweeps (π_SPIM_/π_SLC_GX_> 2.76) and improvement sweeps (π_SLC_GX_/π_SLL_> 5.38), respectively. (**c**) Comparison of genes within the putative domestication and improvement sweeps in our study with those reported in Lin et al.^30^. GDS_previous and GDS_current: genes in domestication sweeps identified in Lin et al. and in this study, respectively. GIS_previous and GIS_current: genes in improvement sweeps identified in Lin et al. and in this study, respectively. (**d**) Five GWAS association loci that overlapped with both domestication and improvement sweeps are shown. The Bonferroni significance threshold (2.4×10^−7^) is indicated by the red horizontal dashed lines.

To determine the genetic and phenotypic basis of tomato breeding, we compared selective sweeps with the 129 GWAS loci identified in our study (**Supplementary Table 4**), and observed that 51 out of 129 (39.5%) GWAS loci overlapped with selective sweeps, including three overlapping only with domestication sweeps, 43 only with improvement sweeps and five with both (**Fig. 5d** and **Supplementary Table 12**). These 51 loci corresponded to 92.6% of traits (25 out of 27, except SD and STIL) investigated in our study, suggesting that most yield-related traits have been under artificial selection, especially during tomato improvement, consistent with the phenotypic difference of most traits among SPIM, SLC and SLL (**Supplementary Fig. 4**). Moreover, five loci associated with SSR, FB, FSIN, and IDM were selected during both tomato domestication and improvement, indicating a continuous selection of their corresponding agronomic traits (**Fig. 5d**).

## DISCUSSION

Dissection of the genetic architecture underlying complex agronomic traits among a large number of tomato accessions is helpful to improve the utilization of these germplasms, and provides a foundation for marker-assisted selection in tomato breeding programs. In this study, we evaluated 27 agronomic traits in a core collection of 605 tomato accessions. High heritability of these traits suggested that they are mainly regulated by genetic factors, and abundant variation in this core collection makes it suitable for GWAS (**Supplementary Table 2**). To meet human interest, the domestication of crops is mainly towards higher yields^48^. Significant variation for each of these traits was observed between different groups, suggesting most of these yield-related traits were selected during tomato domestication and improvement (**Supplementary Fig. 4**), which was further supported by the selective sweep analysis (**Supplementary Table 12**).

Association mapping has identified candidate trait-associated genes in various crops such as *Arabidopsis*^49^, rice^50^, maize^51^, peach^23^, watermelon^52^ and tomato^31^. In this study, we identified a total of 239 significant signals corresponding to 129 genome loci associated with the 27 assessed agronomic traits. Of these, many significant signals for organ size, organ number and organ location traits were co-located with candidate genes identified by a previous linkage analysis^28^ (**Supplementary Tables 4** and **5**). In addition, many novel candidate genes were also identified by our GWAS. Furthermore, we successfully mapped one candidate gene (*SlALMT15*) for stomatal density and the other two candidate genes (*SlELF3* and *SlBOP4*) for five traits (FSIN, IL, PL, IDM and STAL) (**Figs. 3** and **4**). On the other hand, GWAS results for stigma exsertion were not consistent with those obtained through QTL mapping^15, 53^. This inconsistency might be due to the polymorphism of *Style2*.*1* was found only between wild and cultivated tomatoes, and *Style2*.*1* genotype has been fixed in cultivated accessions. However, the variation of *Style3* (SL2.50ch01_84029382) is more abundant in cultivated tomatoes and thus easier to be detected by GWAS (**Supplementary Fig. 36** and **Supplementary Table 6**).

In this study, several candidate genes were homologous to those previously identified in *Arabidopsis* and tomato (**Supplementary Table 5**). We found that CLV3/EMBRYO-SURROUNDING REGION genes *CLE2* and *CLE6* were significantly associated with the petal number. It has been reported that the CLE gene family plays important roles in cell-to-cell communication to control the balance between stem cell proliferation and differentiation in plant development^36^. An acyltransferase gene (*Solyc02g081740*) was identified to be associated with FRNS, OTD, OTLD and SN. Recently, an acyl-CoA N-acyltransferase, MND1, was reported to regulate meristem phase change and plant architecture in barley^54^. In this study, several LOB domain genes were identified as candidates to regulate various agronomic traits in tomato, similar to those reported previously^55^.

Our study provides new phenotypic and genetic insights into the variation of yield-related traits in tomato. It also provides a powerful resource for genetic improvement of tomato and other *Solanaceae* crops. Genes and the possible causative SNPs identified here could be used as potential targets for marker-assistant breeding and/or engineering of tomato with enhanced yield and potential stress resistance.

## METHODS

### Primers

Primers used in this study are listed in **Supplementary Table 13**.

### Plant materials and sequencing

We used a diverse worldwide collection of 605 tomato accessions, including 54 *S. pimpinellifolium*, 140 *S. lycopersicum* var *cerasiforme*, 333 *S. lycopersicum*, 66 Guanxi (GX), 2 *S. habrochaites*, 3 *S. cheesmaniae*, 1 *S. neorickii*, 4 *S. peruvianum*, 1 *S. galapagense* and 1 *S. corneliomuelleri*, for phenotypic and genotypic survey and GWAS analysis. The 66 GX accessions were collected from northwest mountainous region of Guangxi, China, while the remaining 539 accessions and their genotypes were obtained from previous studies^30, 31, 32^. Genomic DNA of the 66 GX accessions was isolated from young leaves using the CTAB method^56^. Paired-end libraries with insert sizes of ∼450-500 bp were constructed using Illumina TruSeq DNA Sample Prep kit according to the manufacturer’s instructions, and sequenced on the Illumina HiSeq 2000 platform. Raw reads were processed to remove adaptor and low-quality sequences (base quality of more than 50% bases ≤5), yielding 7.6-10.9 Gb sequences for each of the 66 GX accessions. All library construction, sequencing and sequence processing were carried out by BGI-Shenzhen, China.

### Phenotyping

Three biological replicates of the entire population were first grown in 15 greenhouses at the Wuhan Academy of Agricultural Sciences, China. Seeds were sown in 50-hole trays containing the soil, peat, and vermiculite (1:1:1) in early March 2016 and then transplanted to the field in mid-April 2016. For each replicate, the accessions were grown in a randomized design, with 12 plants for each accession. Field management, including irrigation, fertilizer application and pest control, followed essentially normal agricultural practice.

The 27 agronomic traits for association analysis were classified into three categories: organ location traits (FIN, FSIN, IL and IDM), organ number traits (FB, FLNS, FLNT, FRNS, FRNT, IT, PN, SD and SN), and organ size traits (FL, FSD, FSL, OLD, OTD, OTLD, PL, SE, SL, SPR, SS, SSR, STAL and STIL). All four organ location traits, all nine organ number traits and three organ size traits (FL, SE and SS) were evaluated by manual observation and counting. The eight organ size traits (FSD, FSL, OLD, OTD, PL, SL, STAL and STIL) were analyzed by vernier caliper in ten fruits or flowers, which were randomly picked from six plants for each accession in each biological replicate. OTLD was calculated as OTD/OLD, SPR was calculated as SL/PL and SSR was calculated as STAL/(STIL+OLD) (**Supplementary Note**).

For stomatal density investigation, seeds were sown in 50-hole trays filled with a 1:1:1 mixture of peat, vermiculite and soil. Subsequently, seedlings were grown at 20/28 °C under a 16 h day/8 h night photoperiod of natural light in a greenhouse. The third fully expanded leaves from the top were sampled after 5-weeks of grown. The lower epidermal strips were peeled and stained with toluidine blue and then loaded onto slides. The number of stomata was observed and counted under an Olympus BHS/BHT microscope (BH-2) at a field of 200×: ≈0.065mm^2^.

### Phenotypic data processing and statistical analysis

For phenotypic values of each agronomic trait, frequency distribution analysis was preformed using EXCEL 2010. The coefficient of variation was calculated independently for each agronomic trait as σ/μ, where σ and μ are the standard deviation and mean of each agronomic trait in the association panel, respectively. Phenotypic data collected from the three biological sample sets were used to calculate broad-sense heritability (*H*^*2*^) as previously described^33^. Significance analysis of difference in the agronomic traits among the three subgroups (SPIM, SLC and SLL) were conducted using the ANOVA test.

### SNP identification and GWAS

The cleaned paired-end reads of the 66 GX accessions were aligned to the tomato reference genome^1^ (Heinz 1706; version SL2.5) using BWA^57^ with default parameters. The SNP dataset of 539 previously reported tomato accessions were download from the Sol Genomics Network (https://solgenomics.net/). The SNP calling of all the 605 accessions was performed by adding the allele information of GX accessions to the SNP dataset of 539 accessions using GATK^58^. Only SNPs with the minor allele frequency (MAF) ≥ 0.05 and the minor allele present in at least six accessions were kept and further used to perform the genome-wide association study.

We used a compressed mixed linear model^59^ to detect the associations between SNPs and the 27 agronomic traits using GEMMA^60^. The genome-wide suggestive and significance thresholds of associations were set at *P*=1/n and *P*=0.05/n, respectively, where n is the effective number of independent SNPs^33^, which corresponded to *P*=2.4×10^−7^ and *P*=1.2×10^−8^. Pairwise LD between the suggestive/significant SNPs for each agronomic trait was calculated using the Plink software (v1.9)^61^. All 239 significant lead SNPs on the tomato genome were integrated into different loci by dividing the whole genome into 200-kb partitions, and the number of significant loci was counted.

### Population and phylogenetic analyses

For the phylogenetic analysis, SNPs of all accessions were first filtered with a missing data rate less than 15% and MAF >0.05. The resulting SNPs at fourfold degenerated sites (23,635) were then used to construct a neighbor-joining tree using the IQTREE software^62^ with 500 bootstrap replicates. PCA was performed with Plink (v1.9)^61^. The population structure of the tomato accessions was inferred using fastStructure^63^ with all SNPs for each *K* (*K*=2-4). Genome-wide π and *F*_ST_ were calculated with VCFtools^64^ (v0.1.15) using 100-kb sliding windows with a step size of 10 kb for each sub-population.

### Identification of domestication and improvement sweeps

For selection sweep analysis, we combined SLC and GX groups into a single group (SLC_GX) to exclude the potential effect of genetic drift. To identify genomic regions affected by domestication and improvement, we first measured the level of genetic diversity (π) within 100-kb sliding windows with a step size of 10 kb in SPIM, SLC_GX and SLL. Candidate domestication and improvement sweeps were identified with the top 5% largest π_SPIM_/π_SLC_GX_ (2.76) and π_SLC_GX_/π_SLL_ (5.33) values, respectively. Finally, windows that were ≤100 kb apart were merged into a single selected region.

### Candidate gene sequencing

To identify the genotype of SL2.50ch03_60427735 in the tomato population, DNA fragments containing this SNP were amplified by PCR in 146 tomato accessions. To detect the variation in the *SlALMT15* gene region, DNA sequences of *SlALMT15* in 13 tomato accessions (TS-9, TS-53, TS-67, TS-91, TS-572, TS-577, TS-604 with high-density stomata; TS-52, TS-55, TS-210, TS-531, np3, and db6 with low-density stomata) were amplified by PCR using primers listed in **Supplementary Table 13**. The PCR products were sequenced and compared against the reference genome for polymorphism analysis.

### RNA isolation and expression analysis

Total RNA was isolated from various tissue of Ts-9, Ts-52, Ts-53 and Ts-55 using TRIZOL reagent (Invitrogen, USA). cDNAs were synthesized from the total RNA using HiScript®II Reverse Transcriptase (Vazyme, Miramar Beach, FL, USA), according to the manufacturer’s protocol. Gene expression was quantified using qRT-PCR as previously described^65^ using primer pairs listed in **Supplementary Table 13**. The *Actin* gene (*Solyc11g008430*) was used as an internal standard. The relative expression of specific genes was quantified using the comparative *C*_T_ method.

### CRISPR/Cas9 construct design and transformation

We used CRISPR/Cas9 binary vectors^66^ (pTX), in which the target sequence was driven by the tomato U6 promoter and Cas9 by 2×35S, for editing of the *SlALMT15* gene. Two *SlALMT15*-specific target sites (sgRNA1 and sgRNA2) were manually selected. The recombinant pTX vector was designed to produce defined deletions within the coding sequence of *SlALMT15* using two sgRNAs alongside the Cas9 endonuclease gene. The plasmid with the correct sgRNA insertion was introduced into *Agrobacterium tumefaciens* strain C58 by electroporation and subsequently transformed into the tomato genome via explants of cotyledon. The high-density stomata accession TS-9 (Ailsa Craig) was used for transformation. Positive detection of T_0_ plants was conducted by PCR using Cas9-specific primers. The CRISPR/Cas9-induced mutations were further genotyped by PCR sequencing using CRISPR/Cas9 detection primers.

### Drought tolerance assays and measurement of physiological indexes

The T_2_ *SlALMT15* knockout lines and wild-type plants were grown at 28°C/20°C (day/night) with a 16 h photoperiod in 10 cm (diameter) plastic pots in a greenhouse with a medium light intensity (approximately 160 μmoles of photons m^-2^ s^-1^). For drought tolerance testing, five-week-old seedling plants were fully watered. Water was then withheld from the seedlings in drought treatment group for 8 d. The control group of seedlings were watered every two days. The phenotype was investigated and recorded at the end of the drought treatment. The third fully expanded leaves from the control and stressed plants were used for the determination of physiological indicators.

The net photosynthetic rate, transpiration rate and stomatal conductance were measured using the LI-6400XT Portable Photosynthesis System (LI-COR, USA) following the manufacturer’s instructions. The values of photosynthesis at 400 μmol·mol-1 minus the values of photosynthesis at 0 μmol·mol-1 were used to approximately represent the photosynthesis capacity^67^. At least five replicated plants were used for each measurement.

The MDA levels were measured as previously described^68^. Briefly, 200 mg of ground leaf samples was homogenized using 3 mL of 5% tri-chloroacetic acid (TCA) and incubated for 20 min. After centrifuged at 5500 rpm for 25 min, 2 mL of supernatant was mixed with 2 mL of 0.67% thiobarbituric acid (TBA) in 10% TCA. After 30 min incubating in boiling water, the mixture was centrifuged and the absorbance of the supernatant was determined spectrophotometrically at 450, 532 and 600 nm. The content of MDA (μmol/g) was calculated as (6.45×(A_532_-A_600_)-0.56×A_450_×V)/(1000×W), where V represents the volume of the extraction buffer (mL), and W represents the weight of the sample (g).

## Data availability

Raw sequences of the 66 GX accessions have been deposited in the Sequence Read Archive (SRA) of National Center for Biotechnology Information under the PRJNA666021. The SNPs have also been deposited into Figshare database (https://figshare.com/collections/_/5136047).

## Acknowledgements

This work was supported by grants from the National Natural Science Foundation of China (31872118 to ZY, 31801861 to JY), China Postdoctoral Science Foundation (2016M592343 to JY) and the US National Science Foundation (IOS-1855585 to Z.F. and J.J.G.).

## Author contributions

Z.Y., Z.F., and J.Y. designed the experiments and managed the project. H.L., J.J.G., X.W., Y.Z., B.O., and J.Z. contributed to the original concept of the project. J.Y., W.W., Y. W., and P.S. collected samples and performed phenotyping. J.Y., X.W., H.Y., G.A. and C.L. performed the data analyses. J.Y. wrote the manuscript. Z.Y., Z.F., and X.W. revised the manuscript.

## Competing interests

The authors declare no competing interests.

## Notes

### Competing Interest Statement

The authors have declared no competing interest.

