## Supplementary Figures for "Genome-wide association study reveals the genetic architecture of 27 yield-related traits in tomato"

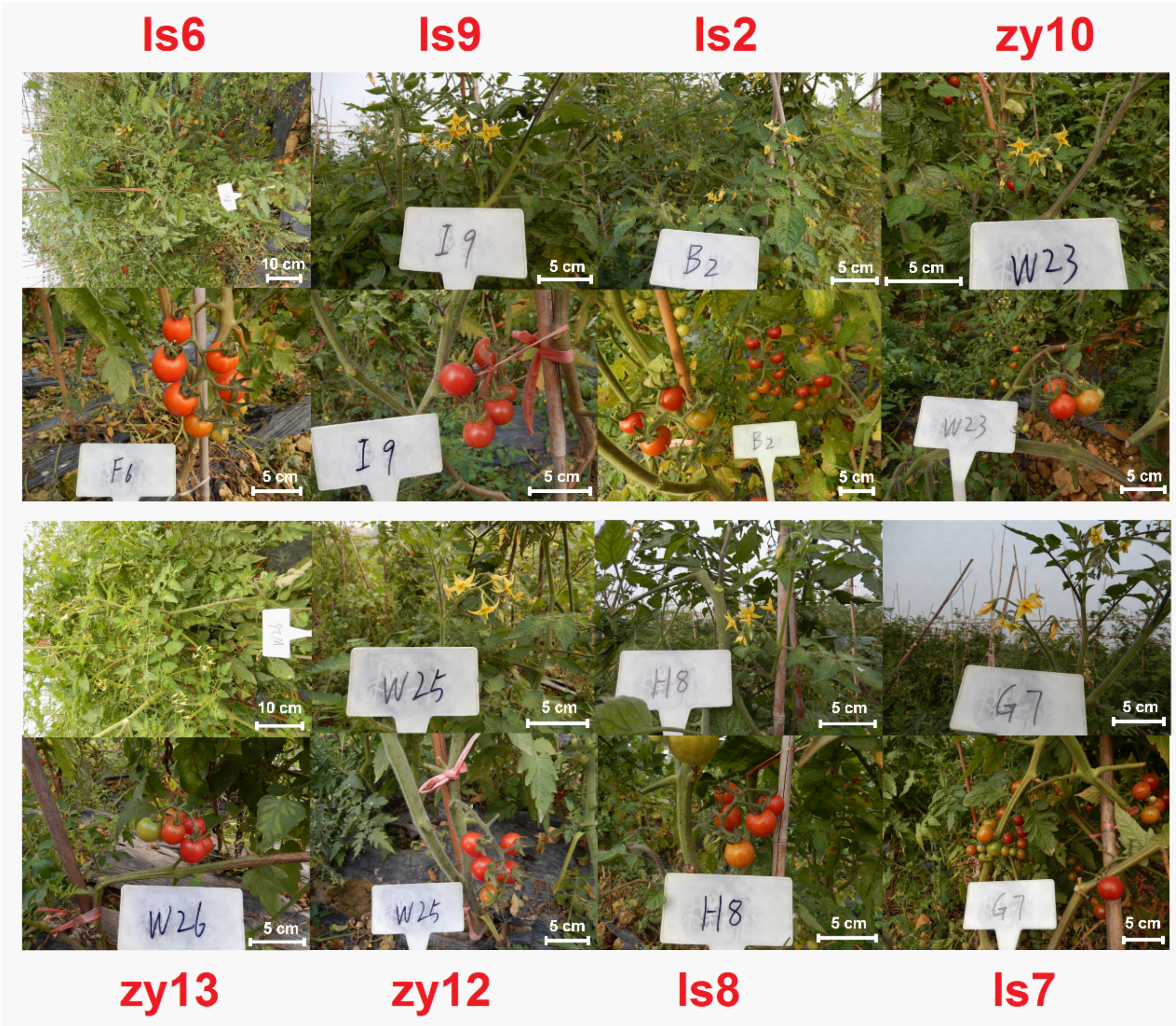


**Supplementary Figure 1** Field phenotypes of some Guangxi tomatoes.

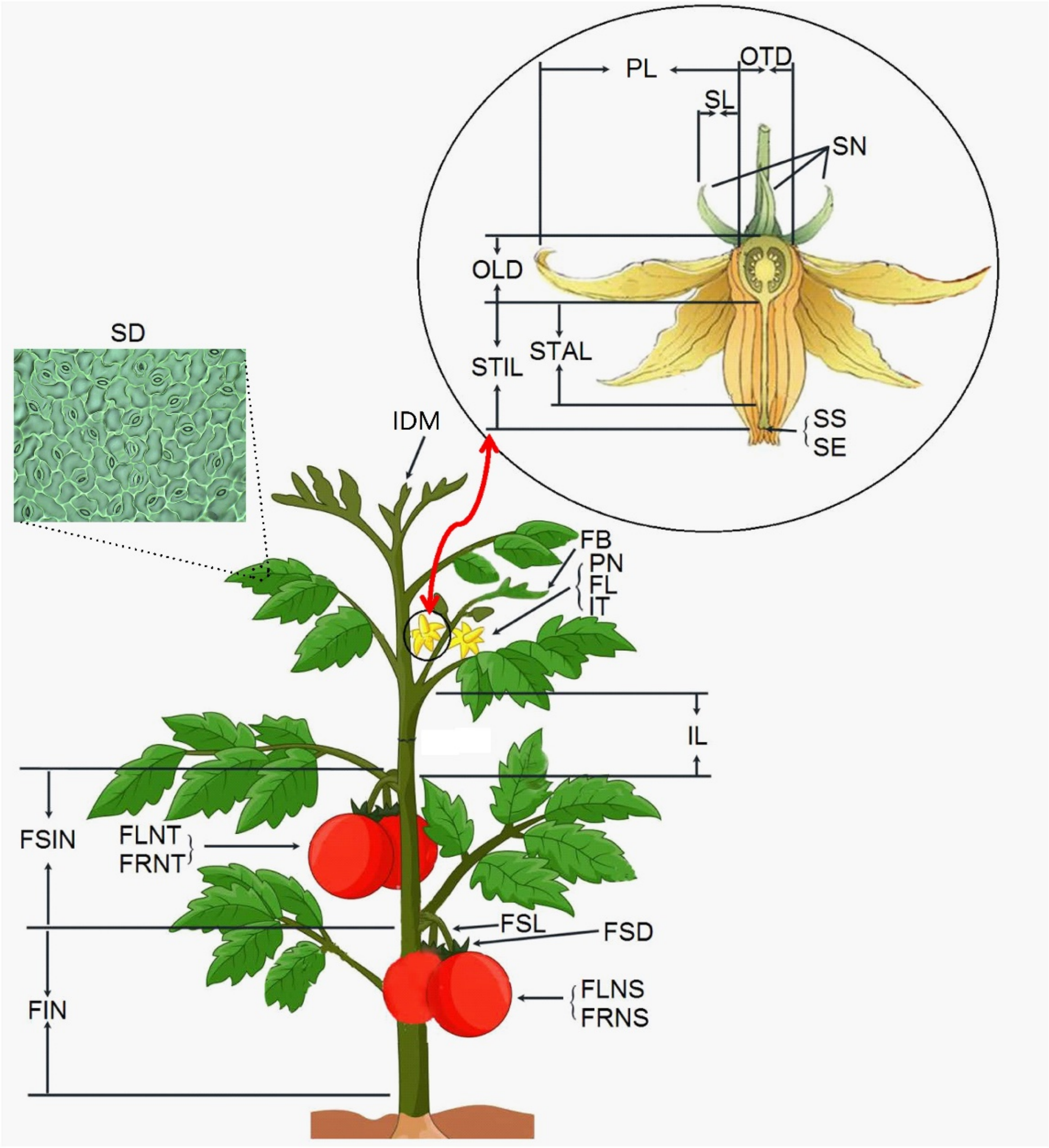


**Supplementary Figure 2 Diagram of the measured 27 agronomic traits in our study**: FIN (first inflorescence node), FSIN (first to second inflorescence node), IL (internode length), IDM (indeterminate or determinate meristem), FB (flower branch), FLNS (flower number on the second inflorescence), FLNT (flower number on the third inflorescence), FRNS (fruit number on the second truss), FRNT (fruit number on the third truss), IT (inflorescence type), PN (petal number), SD (stomatal density), SN (sepal number), FL (fasciated flower), FSD (fruit stalk diameter), FSL (fruit stalk length), OLD (ovary longitudinal diameter), OTD (ovary transverse diameter), OTLD (ovary transverse diameter to ovary longitudinal diameter ratio), PL (petal length), SE (stigma exsertion), SL (sepal length), SPR (sepal length to petal length ratio), SS (stigma shape), SSR (stamen length to (stigma length+ovary longitudinal diameter) ratio), STAL (stamen length) and STIL (stigma length).

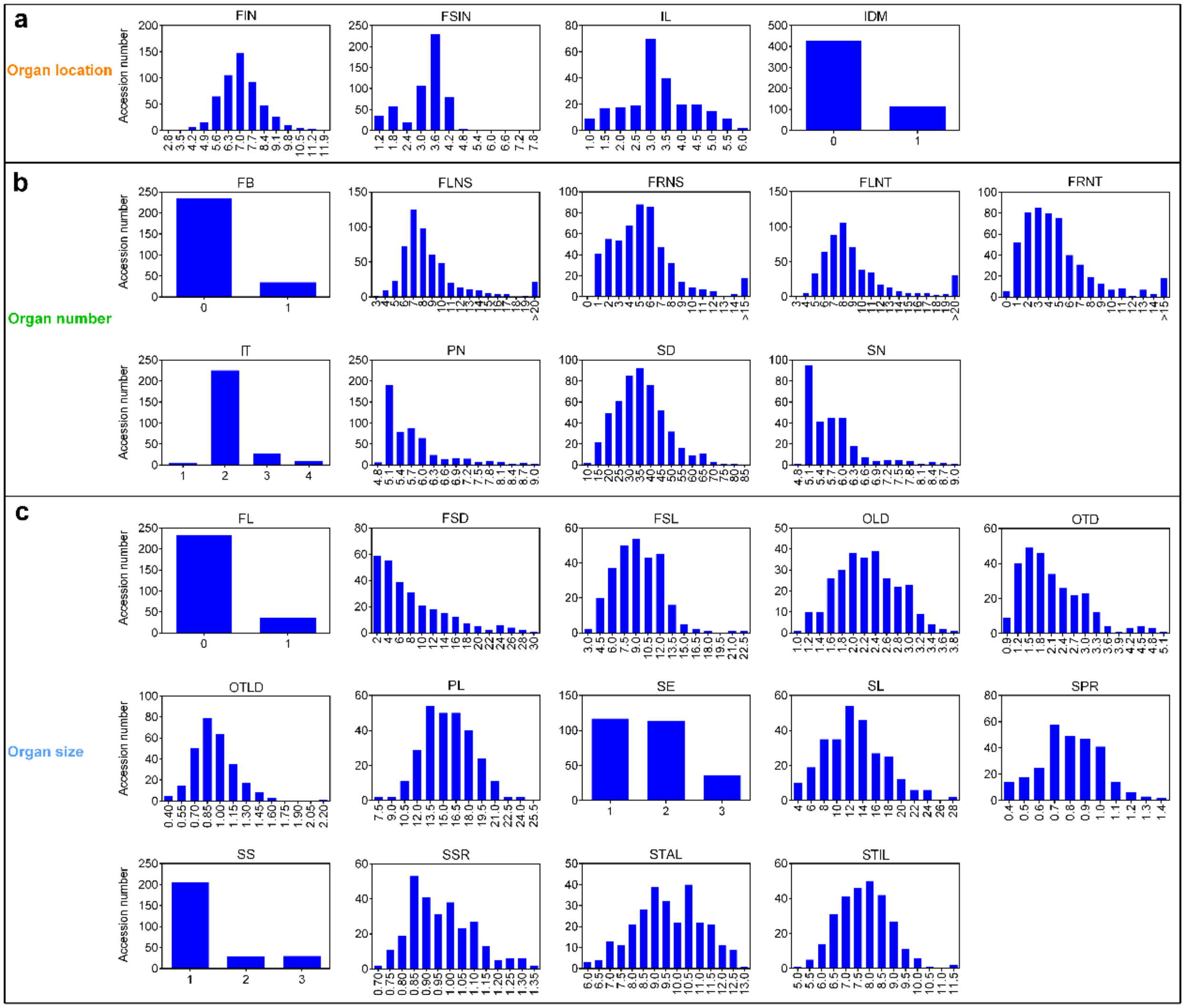


**Supplementary Figure 3: Frequency distribution of phenotypic values of 27 agronomic traits.** The 27 agronomic traits were classified into three categories: 4 organ location traits (**a**), 9 organ number traits (**b**) and 14 organ size traits (**c**). FIN: first inflorescence node, FSIN: first to second inflorescence node, IL: internode length, IDM: indeterminate or determinate meristem, FB: flower branch, FLNS: flower number on the second inflorescence, FLNT: flower number on the third inflorescence, FRNS: fruit number on the second truss, FRNT: fruit number on the third truss, IT: inflorescence type, PN: petal number, SD: stomatal density, SN: sepal number, FL: fasciated flower, FSD: fruit stalk diameter, FSL: fruit stalk length, OLD: ovary longitudinal diameter, OTD: ovary transverse diameter, OTLD: ovary transverse diameter to ovary longitudinal diameter ratio, PL: petal length, SE: stigma exsertion, SL: sepal length, SPR: sepal length to petal length ratio, SS: stigma shape, SSR: stamen length to (stigma length+ovary longitudinal diameter) ratio, STAL: stamen length and STIL: stigma length.

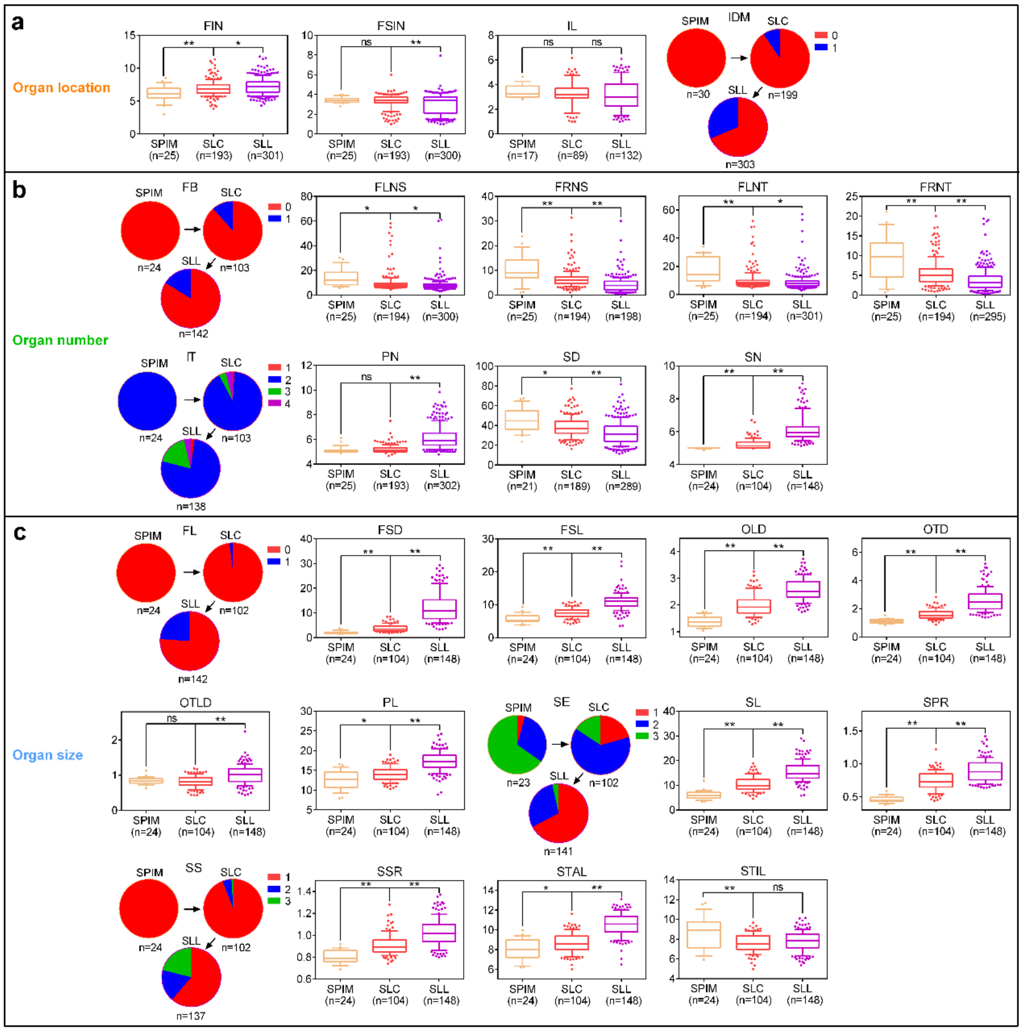


**Supplementary Figure 4: Phenotypic distribution of 27 agronomic traits in the different tomato subgroups. (a)** 4 organ location traits, **(b)** 9 organ number traits, **(c)** 14 organ size traits. SPIM, *S. pimpinellifolium*. SLC, *S. lycopersicum* var. *cerasiforme*. SLL, *Solanum lycopersicum* var. *lycopersicum*. FIN: first inflorescence node, FSIN: first to second inflorescence node, IL: internode length, IDM: indeterminate or determinate meristem, FB: flower branch, FLNS: flower number on the second inflorescence, FLNT: flower number on the third inflorescence, FRNS: fruit number on the second truss, FRNT: fruit number on the third truss, IT: inflorescence type, PN: petal number, SD: stomatal density, SN: sepal number, FL: fasciated flower, FSD: fruit stalk diameter, FSL: fruit stalk length, OLD: ovary longitudinal diameter, OTD: ovary transverse diameter, OTLD: ovary transverse diameter to ovary longitudinal diameter ratio, PL: petal length, SE: stigma exsertion, SL: sepal length, SPR: sepal length to petal length ratio, SS: stigma shape, SSR: stamen length to (stigma length+ovary longitudinal diameter) ratio, STAL: stamen length and STIL: stigma length. For each box plot, the horizontal line in the box indicates the median value, the box height indicates the $25^{th}$ to $75^{th}$ percentile of the total data, the whiskers indicate the interquartile range, and the outer dots indicate outliers. Asterisks indicate significant differences by *t* test: ns, *P*>0.05, *0.05<*P*<0.01, ***P*<0.01.

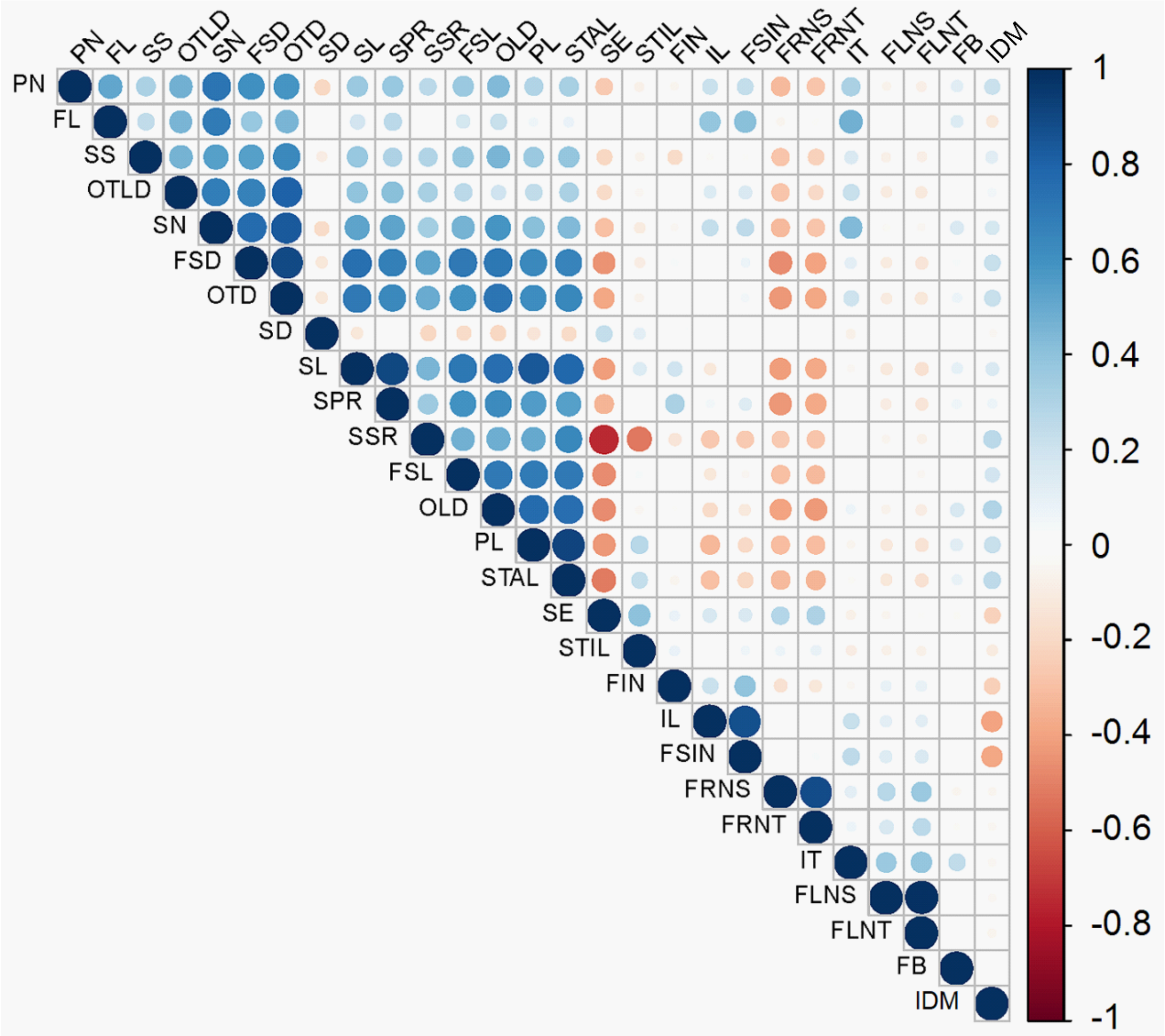


**Supplementary Figure 5: Correlations between the analyzed phenotypes.** The correlation coefficient (Spearman) ranges from -1 (red color) to +1 (blue color). FIN: first inflorescence node, FSIN: first to second inflorescence node, IL: internode length, IDM: indeterminate or determinate meristem, FB: flower branch, FLNS: flower number on the second inflorescence, FLNT: flower number on the third inflorescence, FRNS: fruit number on the second truss, FRNT: fruit number on the third truss, IT: inflorescence type, PN: petal number, SD: stomatal density, SN: sepal number, FL: fasciated flower, FSD: fruit stalk diameter, FSL: fruit stalk length, OLD: ovary longitudinal diameter, OTD: ovary transverse diameter, OTLD: ovary transverse diameter to ovary longitudinal diameter ratio, PL: petal length, SE: stigma exsertion, SL: sepal length, SPR: sepal length to petal length ratio, SS: stigma shape, SSR: stamen length to (stigma length+ovary longitudinal diameter) ratio, STAL: stamen length and STIL: stigma length.

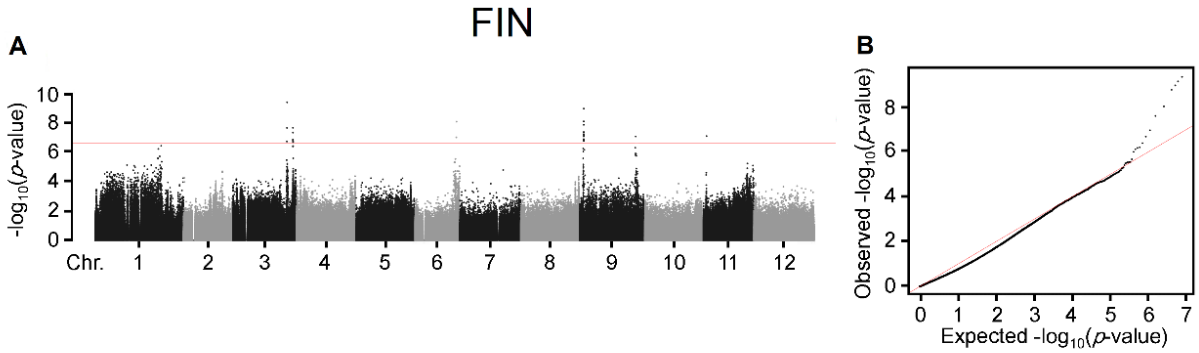


**Supplementary Figure 6: Manhattan plot (a) and Quantile-Quantile (Q-Q) plot (b) of GWAS for FIN (first inflorescence node).** Negative $\log_{10}$-transformed *P* values from the compressed mixed linear model were plotted against SNPs position on each of the 12 chromosomes. The horizontal dashed line indicates a genome-wide significance threshold of $2.4\times10^{-7}$.

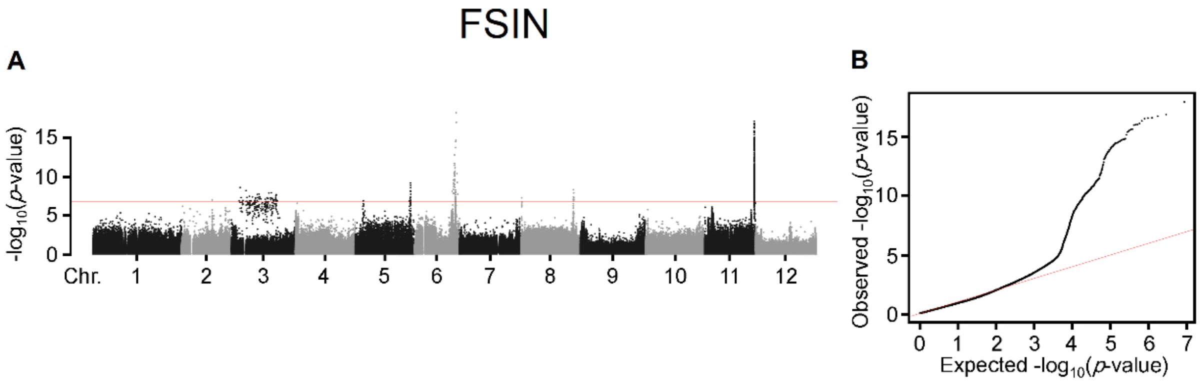


**Supplementary Figure 7: Manhattan plot (a) and Quantile-Quantile (Q-Q) plot (b) of GWAS for FSIN (first to second inflorescence node).** Negative $\log_{10}$-transformed *P* values from the compressed mixed linear model were plotted against SNPs position on each of the 12 chromosomes. The horizontal dashed line indicates a genome-wide significance threshold of $2.4\times10^{-7}$.

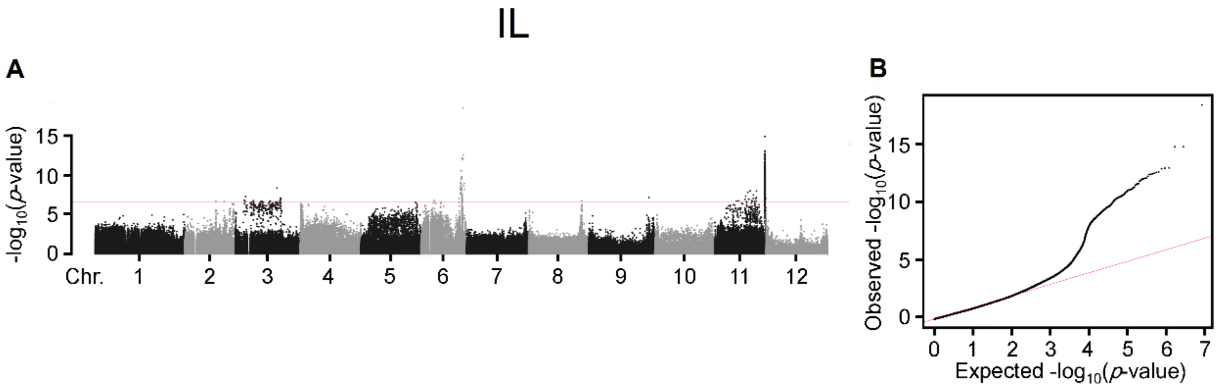


**Supplementary Figure 8: Manhattan plot (a) and Quantile-Quantile (Q-Q) plot (b) of GWAS for IL (internode length).** Negative $\log_{10}$-transformed *P* values from the compressed mixed linear model were plotted against SNPs position on each of the 12 chromosomes. The horizontal dashed line indicates a genome-wide significance threshold of $2.4\times10^{-7}$.

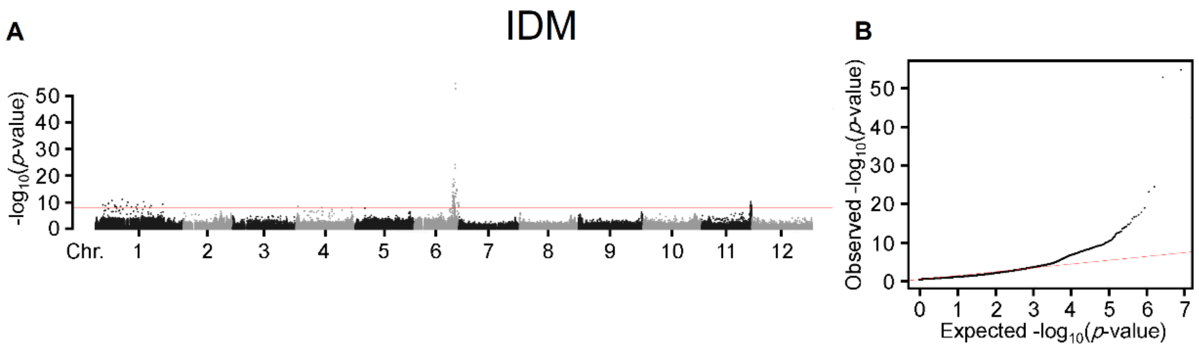


**Supplementary Figure 9: Manhattan plot (a) and Quantile-Quantile (Q-Q) plot (b) of GWAS for IDM (indeterminate or determinate meristem).** Negative $\log_{10}$-transformed *P* values from the compressed mixed linear model were plotted against SNPs position on each of the 12 chromosomes. The horizontal dashed line indicates a genome-wide significance threshold of $2.4 \times 10^{-7}$.

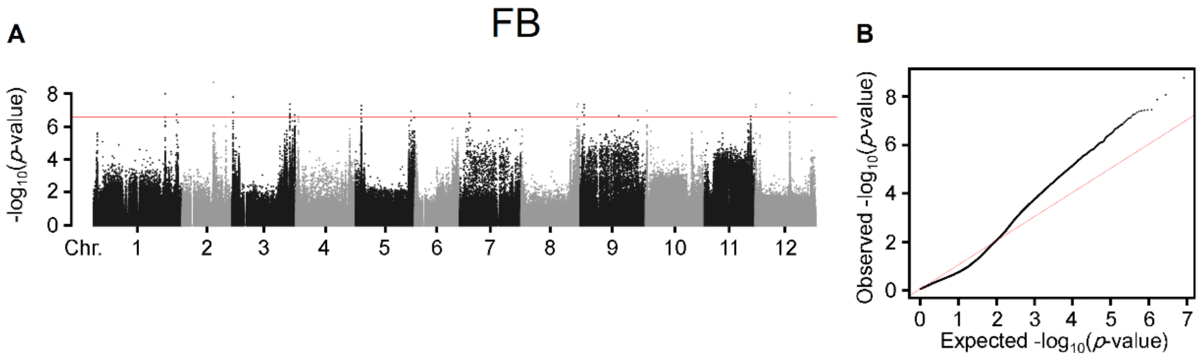


**Supplementary Figure 10: Manhattan plot (a) and Quantile-Quantile (Q-Q) plot (b) of GWAS for FB (flower branch).** Negative $\log_{10}$-transformed *P* values from the compressed mixed linear model were plotted against SNPs position on each of the 12 chromosomes. The horizontal dashed line indicates a genome-wide significance threshold of $2.4\times10^{-7}$.

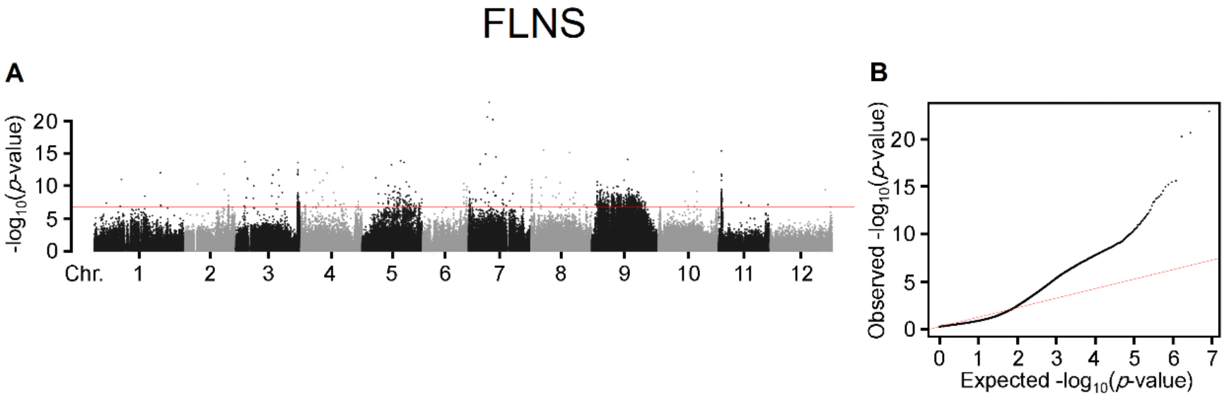


**Supplementary Figure 11: Manhattan plot (a) and Quantile-Quantile (Q-Q) plot (b) of GWAS for FLNS (flower number on the second inflorescence).** Negative $log_{10}$-transformed *P* values from the compressed mixed linear model were plotted against SNPs position on each of the 12 chromosomes. The horizontal dashed line indicates a genome-wide significance threshold of $2.4 \times 10^{-7}$.

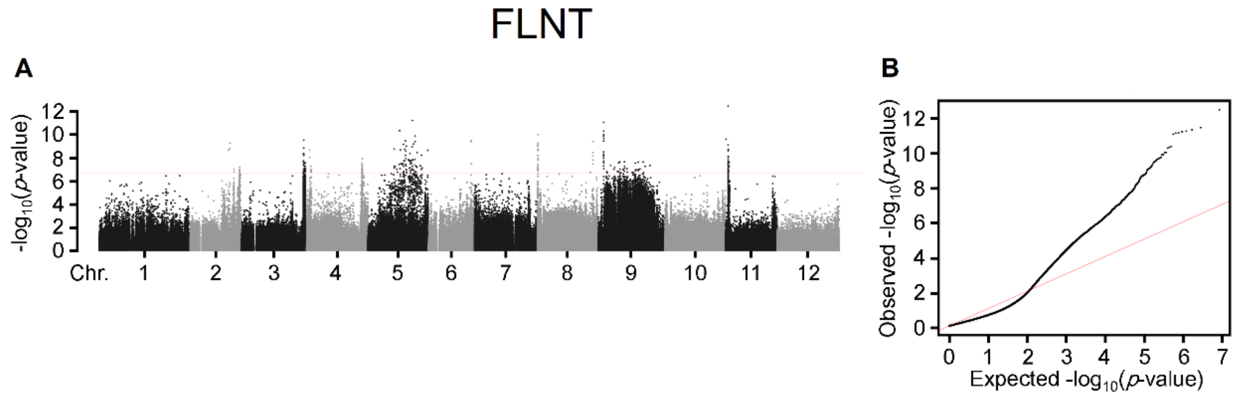


**Supplementary Figure 12: Manhattan plot (a) and Quantile-Quantile (Q-Q) plot (b) of GWAS for FLNT (flower number on the third inflorescence).** Negative $\log_{10}$-transformed *P* values from the compressed mixed linear model were plotted against SNPs position on each of the 12 chromosomes. The horizontal dashed line indicates a genome-wide significance threshold of $2.4\times10^{-7}$.

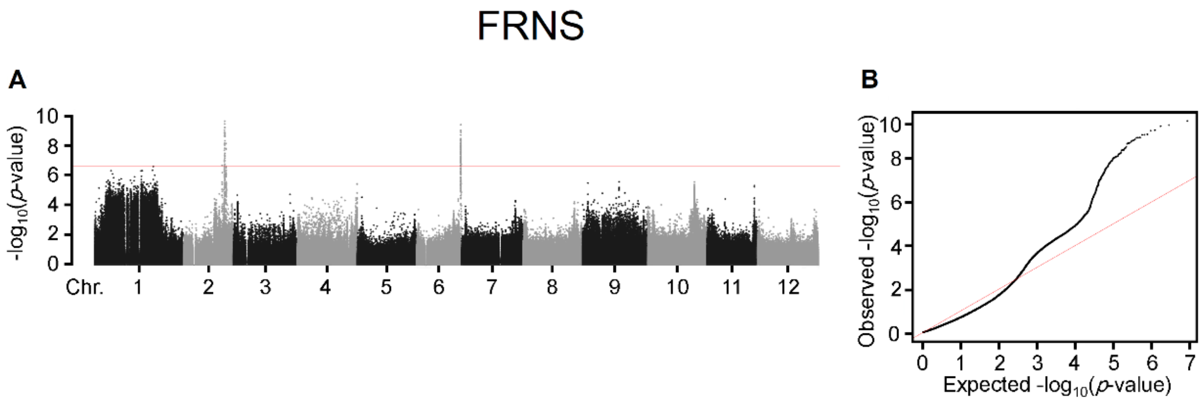


**Supplementary Figure 13: Manhattan plot (a) and Quantile-Quantile (Q-Q) plot (b) of GWAS for FRNS (fruit number on the second truss).** Negative $log_{10}$-transformed *P* values from the compressed mixed linear model were plotted against SNPs position on each of the 12 chromosomes. The horizontal dashed line indicates a genome-wide significance threshold of $2.4{\times}10^{-7}$.

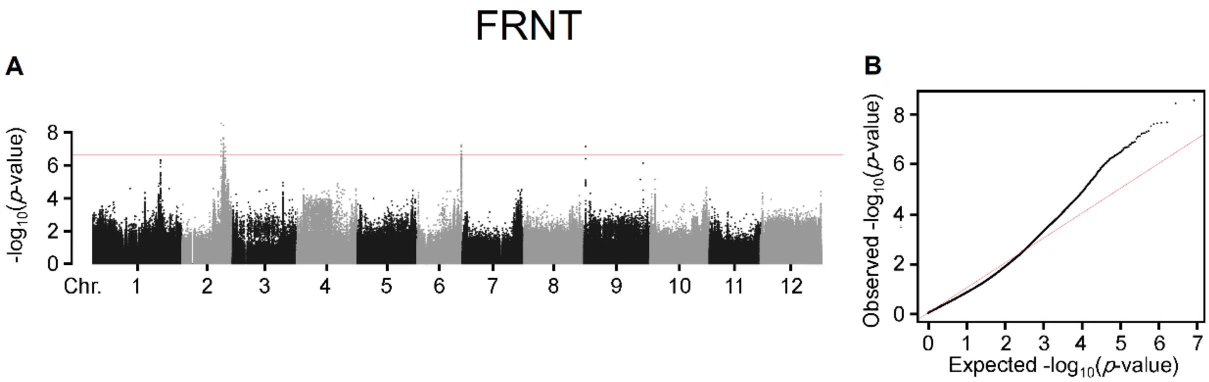


**Supplementary Figure 14: Manhattan plot (a) and Quantile-Quantile (Q-Q) plot (b) of GWAS for FRNT (fruit number on the third truss).** Negative $\log_{10}$-transformed *P* values from the compressed mixed linear model were plotted against SNPs position on each of the 12 chromosomes. The horizontal dashed line indicates a genome-wide significance threshold of $2.4\times10^{-7}$.

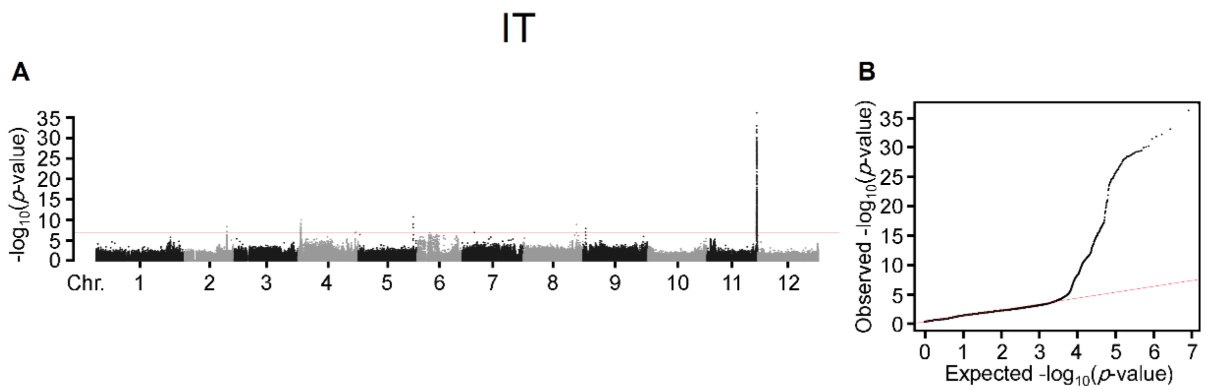


**Supplementary Figure 15: Manhattan plot (a) and Quantile-Quantile (Q-Q) plot (b) of GWAS for IT (inflorescence type).** Negative $log_{10}$-transformed *P* values from the compressed mixed linear model were plotted against SNPs position on each of the 12 chromosomes. The horizontal dashed line indicates a genome-wide significance threshold of $2.4\times10^{-7}$.

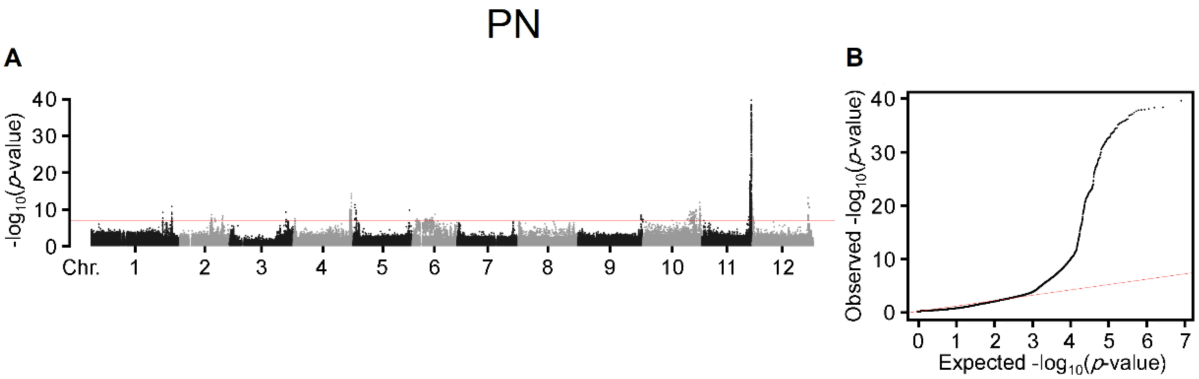


**Supplementary Figure 16: Manhattan plot (a) and Quantile-Quantile (Q-Q) plot (b) of GWAS for PN (petal number).** Negative $\log_{10}$-transformed *P* values from the compressed mixed linear model were plotted against SNPs position on each of the 12 chromosomes. The horizontal dashed line indicates a genome-wide significance threshold of $2.4\times10^{-7}$.

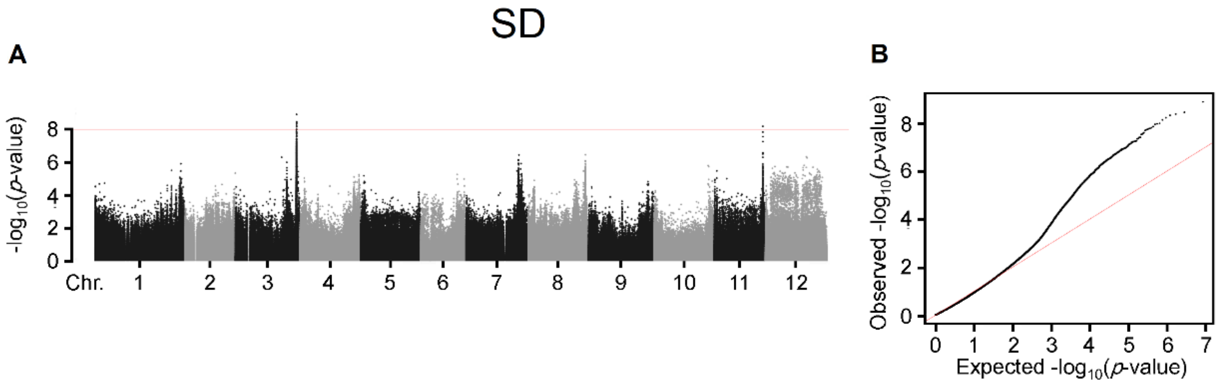


**Supplementary Figure 17: Manhattan plot (a) and Quantile-Quantile (Q-Q) plot (b) of GWAS for SD (stomatal density).** Negative $\log_{10}$-transformed *P* values from the compressed mixed linear model were plotted against SNPs position on each of the 12 chromosomes. The horizontal dashed line indicates a genome-wide significance threshold of $2.4\times10^{-7}$.

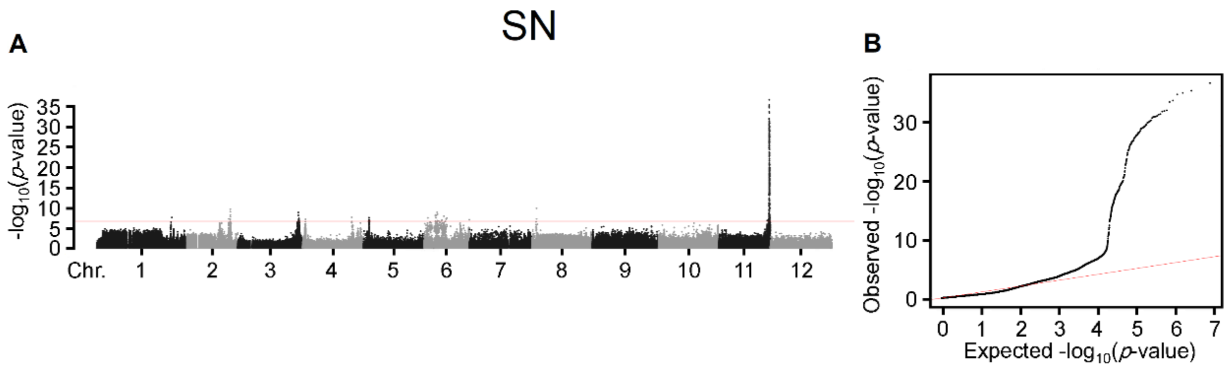


**Supplementary Figure 18: Manhattan plot (a) and Quantile-Quantile (Q-Q) plot (b) of GWAS for SN (sepal number).** Negative $log_{10}$-transformed *P* values from the compressed mixed linear model were plotted against SNPs position on each of the 12 chromosomes. The horizontal dashed line indicates a genome-wide significance threshold of $2.4 \times 10^{-7}$.

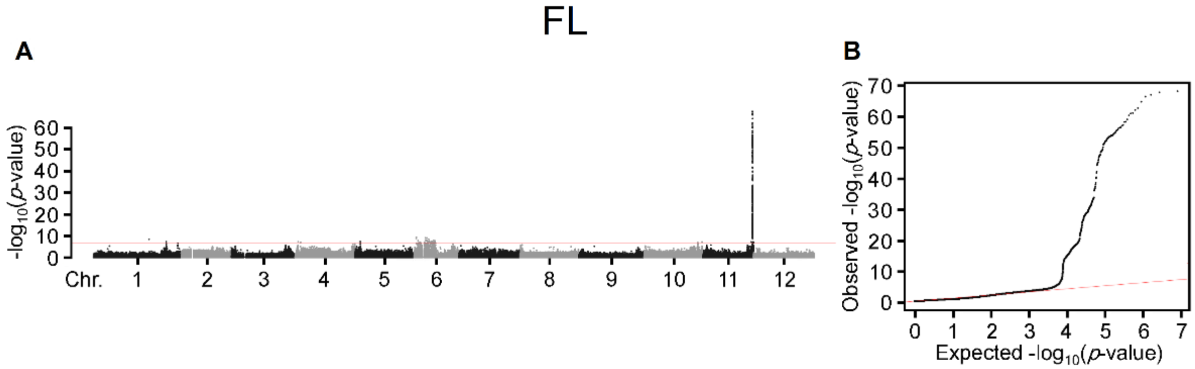


**Supplementary Figure 19: Manhattan plot (a) and Quantile-Quantile (Q-Q) plot (b) of GWAS for FL (fasciated flower).** Negative $log_{10}$-transformed *P* values from the compressed mixed linear model were plotted against SNPs position on each of the 12 chromosomes. The horizontal dashed line indicates a genome-wide significance threshold of $2.4\times10^{-7}$.

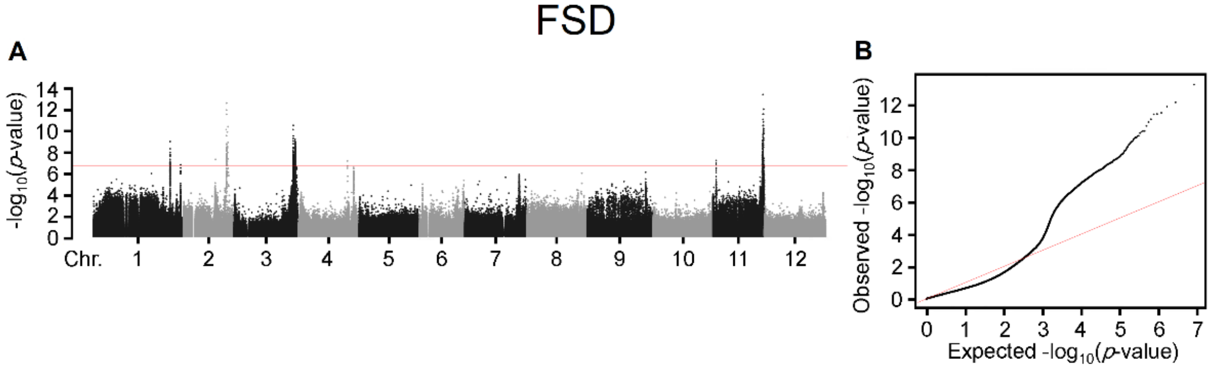


**Supplementary Figure 20: Manhattan plot (a) and Quantile-Quantile (Q-Q) plot (b) of GWAS for FSD (fruit stalk diameter).** Negative $log_{10}$-transformed *P* values from the compressed mixed linear model were plotted against SNPs position on each of the 12 chromosomes. The horizontal dashed line indicates a genome-wide significance threshold of $2.4\times10^{-7}$.

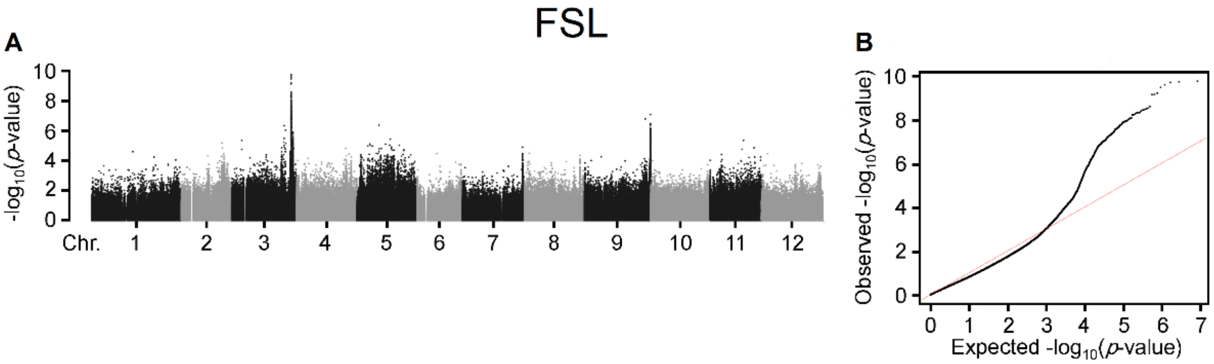


**Supplementary Figure 21: Manhattan plot (a) and Quantile-Quantile (Q-Q) plot (b) of GWAS for FSL (fruit stalk length).** Negative $\log_{10}$-transformed *P* values from the compressed mixed linear model were plotted against SNPs position on each of the 12 chromosomes. The horizontal dashed line indicates a genome-wide significance threshold of $2.4\times10^{-7}$.

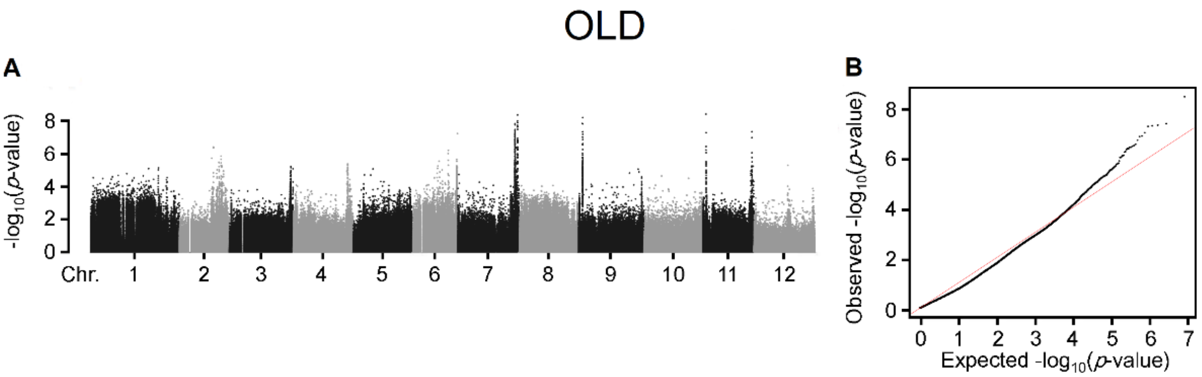


**Supplementary Figure 22: Manhattan plot (a) and Quantile-Quantile (Q-Q) plot (b) of GWAS for OLD (ovary longitudinal diameter).** Negative $\log_{10}$-transformed *P* values from the compressed mixed linear model were plotted against SNPs position on each of the 12 chromosomes. The horizontal dashed line indicates a genome-wide significance threshold of $2.4\times10^{-7}$.

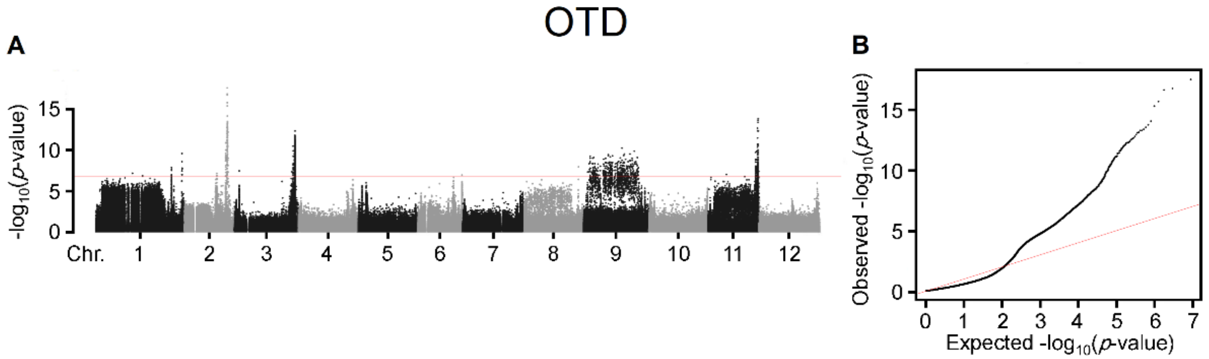


**Supplementary Figure 23: Manhattan plot (a) and Quantile-Quantile (Q-Q) plot (b) of GWAS for OTD (ovary transverse diameter).** Negative $\log_{10}$-transformed *P* values from the compressed mixed linear model were plotted against SNPs position on each of the 12 chromosomes. The horizontal dashed line indicates a genome-wide significance threshold of $2.4\times10^{-7}$.

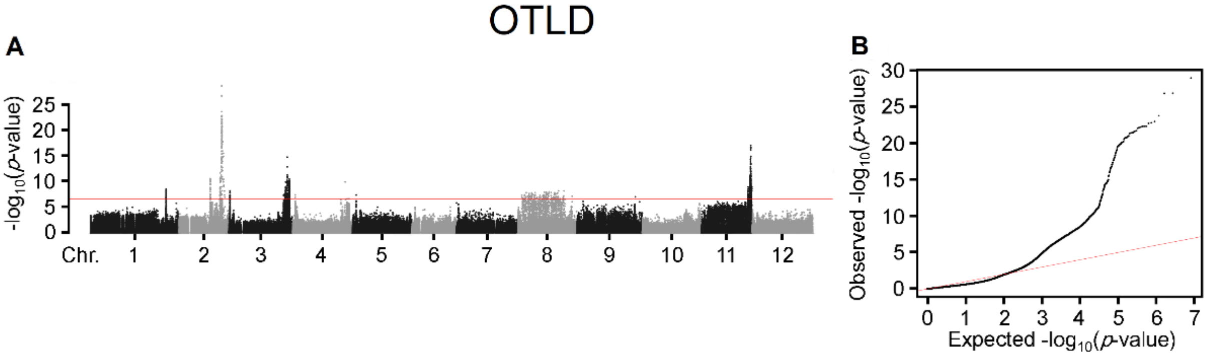


**Supplementary Figure 24: Manhattan plot (a) and Quantile-Quantile (Q-Q) plot (b) of GWAS for OTLD (ovary transverse diameter to ovary longitudinal diameter ratio).** Negative $\log_{10}$-transformed $P$ values from the compressed mixed linear model were plotted against SNPs position on each of the 12 chromosomes. The horizontal dashed line indicates a genome-wide significance threshold of $2.4\times10^{-7}$.

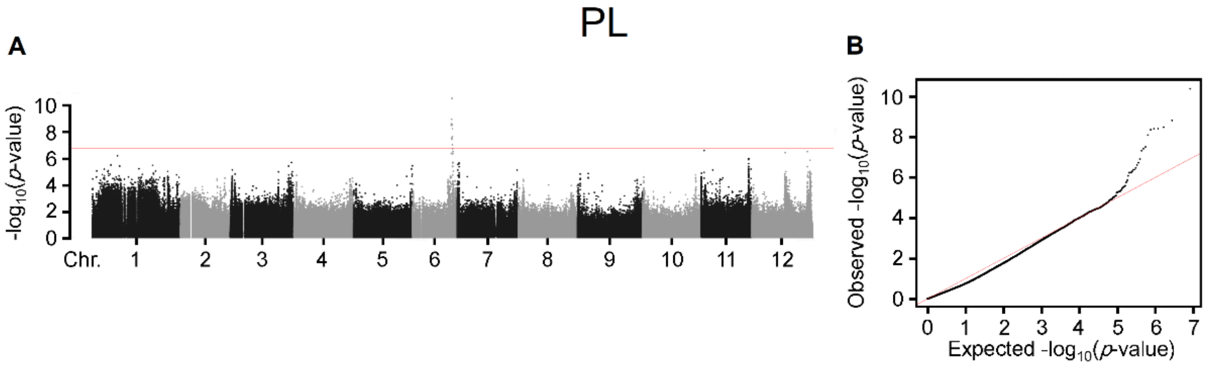


**Supplementary Figure 25: Manhattan plot (a) and Quantile-Quantile (Q-Q) plot (b) of GWAS for PL (petal length).** Negative $\log_{10}$-transformed *P* values from the compressed mixed linear model were plotted against SNPs position on each of the 12 chromosomes. The horizontal dashed line indicates a genome-wide significance threshold of $2.4 \times 10^{-7}$.

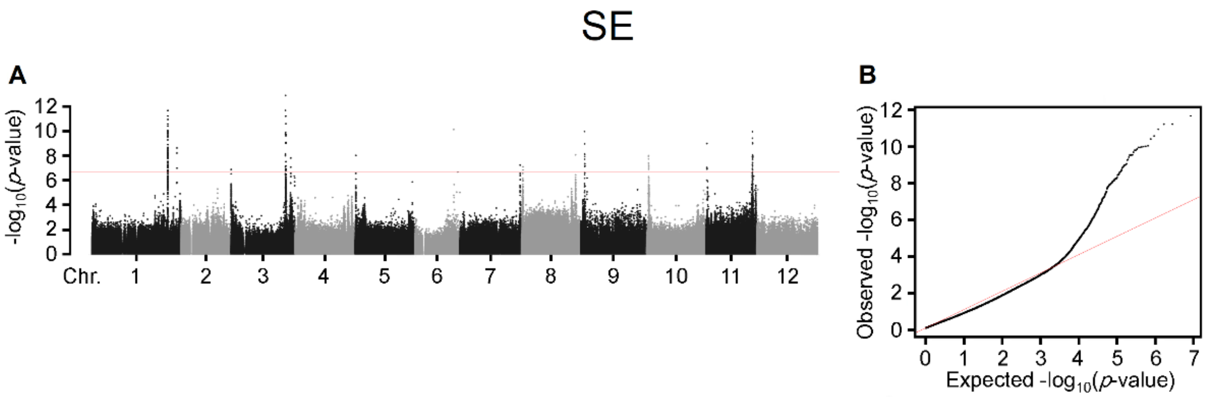


**Supplementary Figure 26: Manhattan plot (a) and Quantile-Quantile (Q-Q) plot (b) of GWAS for SE (stigma exsertion).** Negative $\log_{10}$-transformed *P* values from the compressed mixed linear model were plotted against SNPs position on each of the 12 chromosomes. The horizontal dashed line indicates a genome-wide significance threshold of $2.4 \times 10^{-7}$.

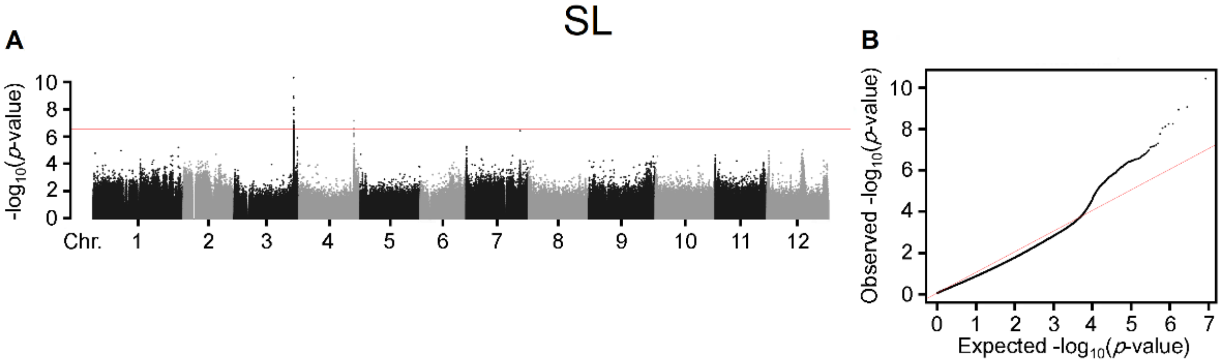


**Supplementary Figure 27: Manhattan plot (a) and Quantile-Quantile (Q-Q) plot (b) of GWAS for SL (sepal length).** Negative $\log_{10}$-transformed *P* values from the compressed mixed linear model were plotted against SNPs position on each of the 12 chromosomes. The horizontal dashed line indicates a genome-wide significance threshold of $2.4\times10^{-7}$.

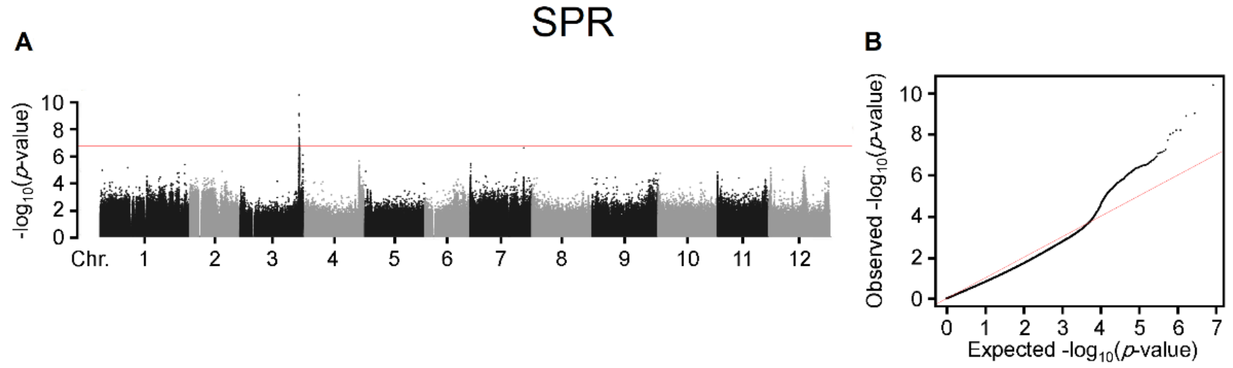


**Supplementary Figure 28: Manhattan plot (a) and Quantile-Quantile (Q-Q) plot (b) of GWAS for SPR (sepal length to petal length ratio).** Negative $\log_{10}$-transformed *P* values from the compressed mixed linear model were plotted against SNPs position on each of the 12 chromosomes. The horizontal dashed line indicates a genome-wide significance threshold of $2.4\times10^{-7}$.

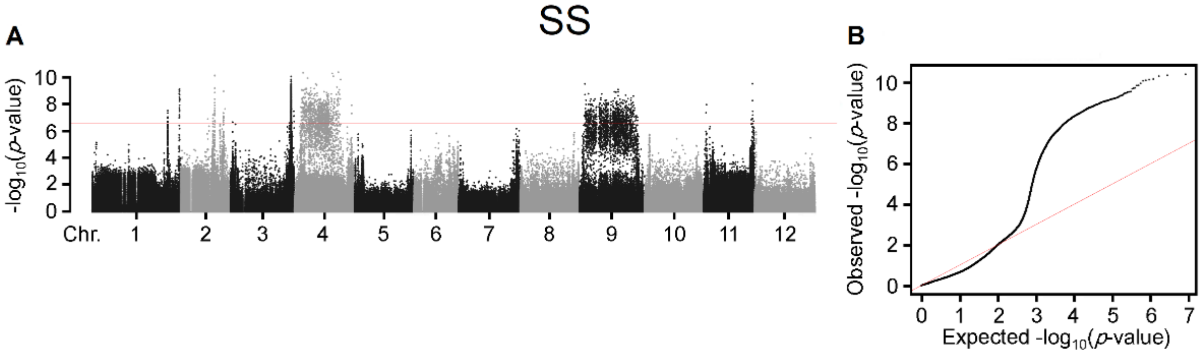


**Supplementary Figure 29: Manhattan plot (a) and Quantile-Quantile (Q-Q) plot (b) of GWAS for SS (stigma shape).** Negative $\log_{10}$-transformed *P* values from the compressed mixed linear model were plotted against SNPs position on each of the 12 chromosomes. The horizontal dashed line indicates a genome-wide significance threshold of $2.4\times10^{-7}$.

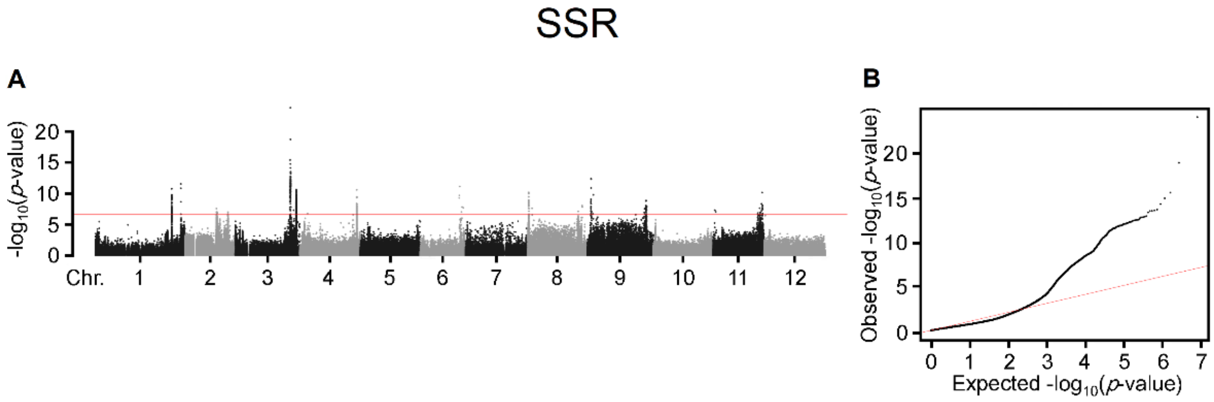


**Supplementary Figure 30: Manhattan plot (a) and Quantile-Quantile (Q-Q) plot (b) of GWAS for SSR (stamen length to (stigma length + ovary longitudinal diameter) ratio).** Negative $\log_{10}$-transformed $P$ values from the compressed mixed linear model were plotted against SNPs position on each of the 12 chromosomes. The horizontal dashed line indicates a genome-wide significance threshold of $2.4\times10^{-7}$.

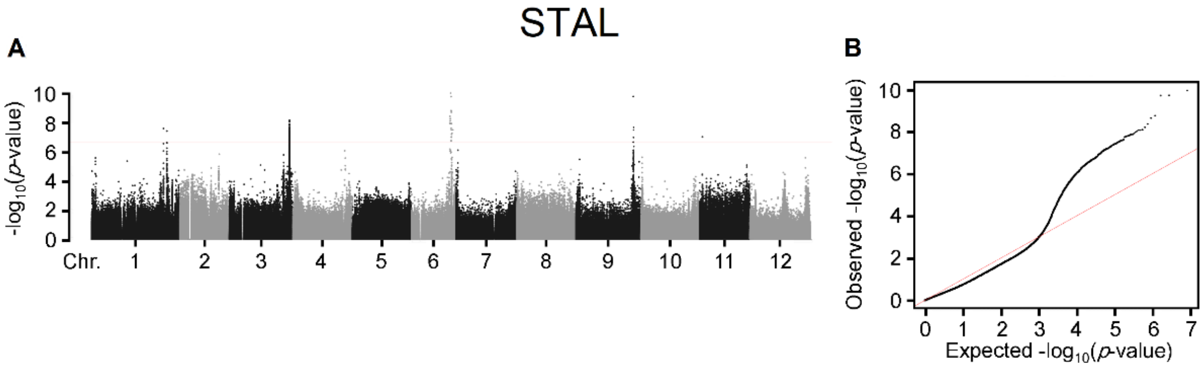


**Supplementary Figure 31: Manhattan plot (a) and Quantile-Quantile (Q-Q) plot (b) of GWAS for STAL (stamen length).** Negative $\log_{10}$-transformed *P* values from the compressed mixed linear model were plotted against SNPs position on each of the 12 chromosomes. The horizontal dashed line indicates a genome-wide significance threshold of $2.4\times10^{-7}$.

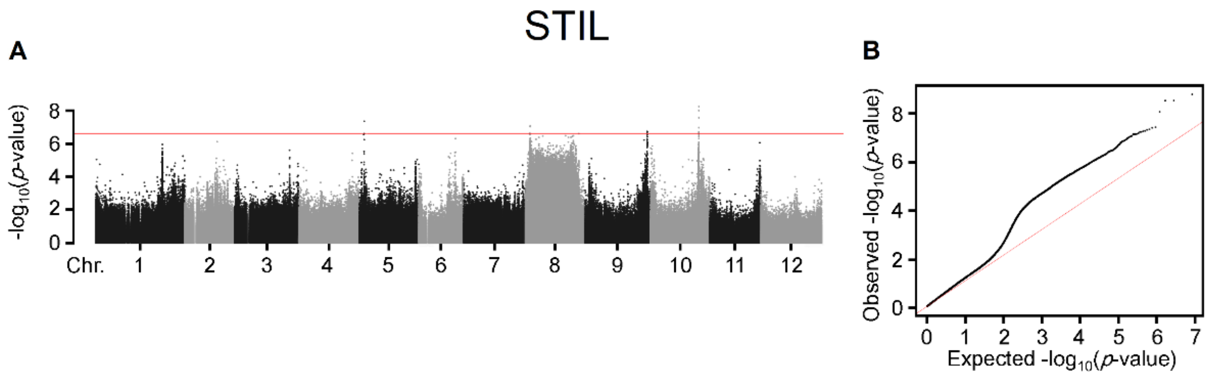


**Supplementary Figure 32: Manhattan plot (a) and Quantile-Quantile (Q-Q) plot (b) of GWAS for STIL (stigma length).** Negative $\log_{10}$-transformed *P* values from the compressed mixed linear model were plotted against SNPs position on each of the 12 chromosomes. The horizontal dashed line indicates a genome-wide significance threshold of $2.4\times10^{-7}$.

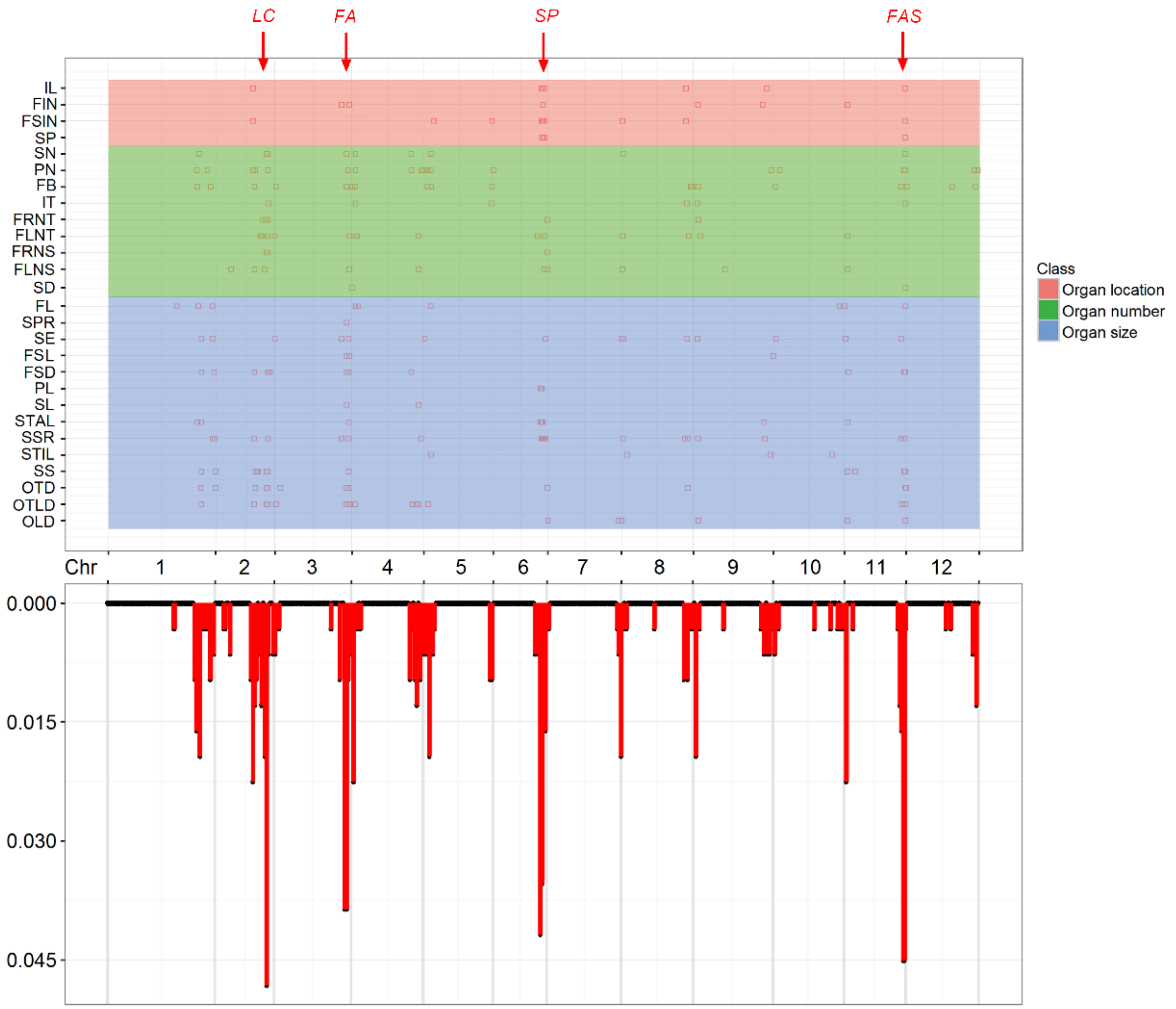


**Supplementary Figure 33: Suggestive loci (*P* < 2.4 × 10$^{-7}$) for the GWAS results associated with 27 tomato traits.** The top figure shows the distribution of the suggestive loci (represented by dots) across the tomato genome. The bottom figure shows the density of the suggestive loci across the tomato genome. Detailed information for all detected loci is provided in Supplementary Table 4. FIN: first inflorescence node, FSIN: first to second inflorescence node, IL: internode length, IDM: indeterminate or determinate meristem, FB: flower branch, FLNS: flower number on the second inflorescence, FLNT: flower number on the third inflorescence, FRNS: fruit number on the second truss, FRNT: fruit number on the third truss, IT: inflorescence type, PN: petal number, SD: stomatal density, SN: sepal number, FL: fasciated flower, FSD: fruit stalk diameter, FSL: fruit stalk length, OLD: ovary longitudinal diameter, OTD: ovary transverse diameter, OTLD: ovary transverse diameter to ovary longitudinal diameter ratio, PL: petal length, SE: stigma exsertion, SL: sepal length, SPR: sepal length to petal length ratio, SS: stigma shape, SSR: stamen length to (stigma length+ovary longitudinal diameter) ratio, STAL: stamen length and STIL: stigma length.

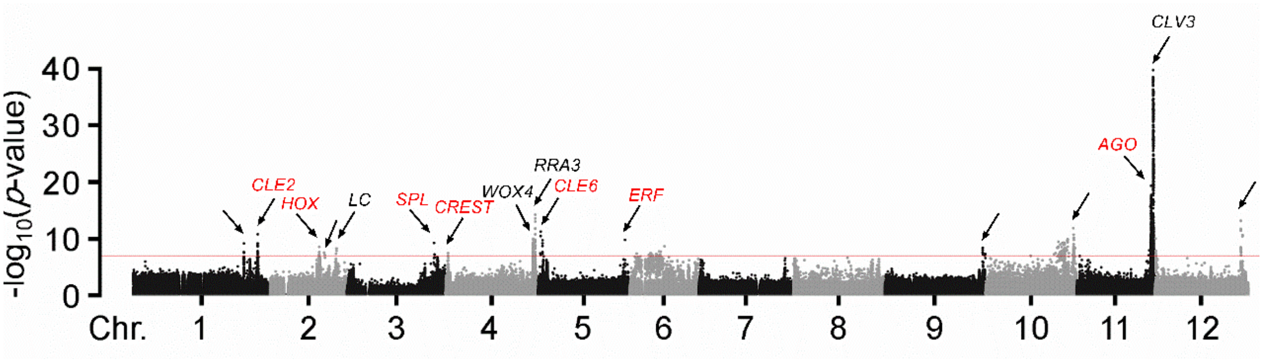


**Supplementary Figure 34: Candidate genes identified by genome-wide association study for PN (petal number).** Negative $\log_{10}$-transformed *P* values from the compressed mixed linear model were plotted against SNPs position on each of the 12 chromosomes. The horizontal dashed line indicates a genome-wide significance threshold of $2.4\times10^{-7}$. The name of known and unknown related genes near the association signals are shown in black and red text, respectively.

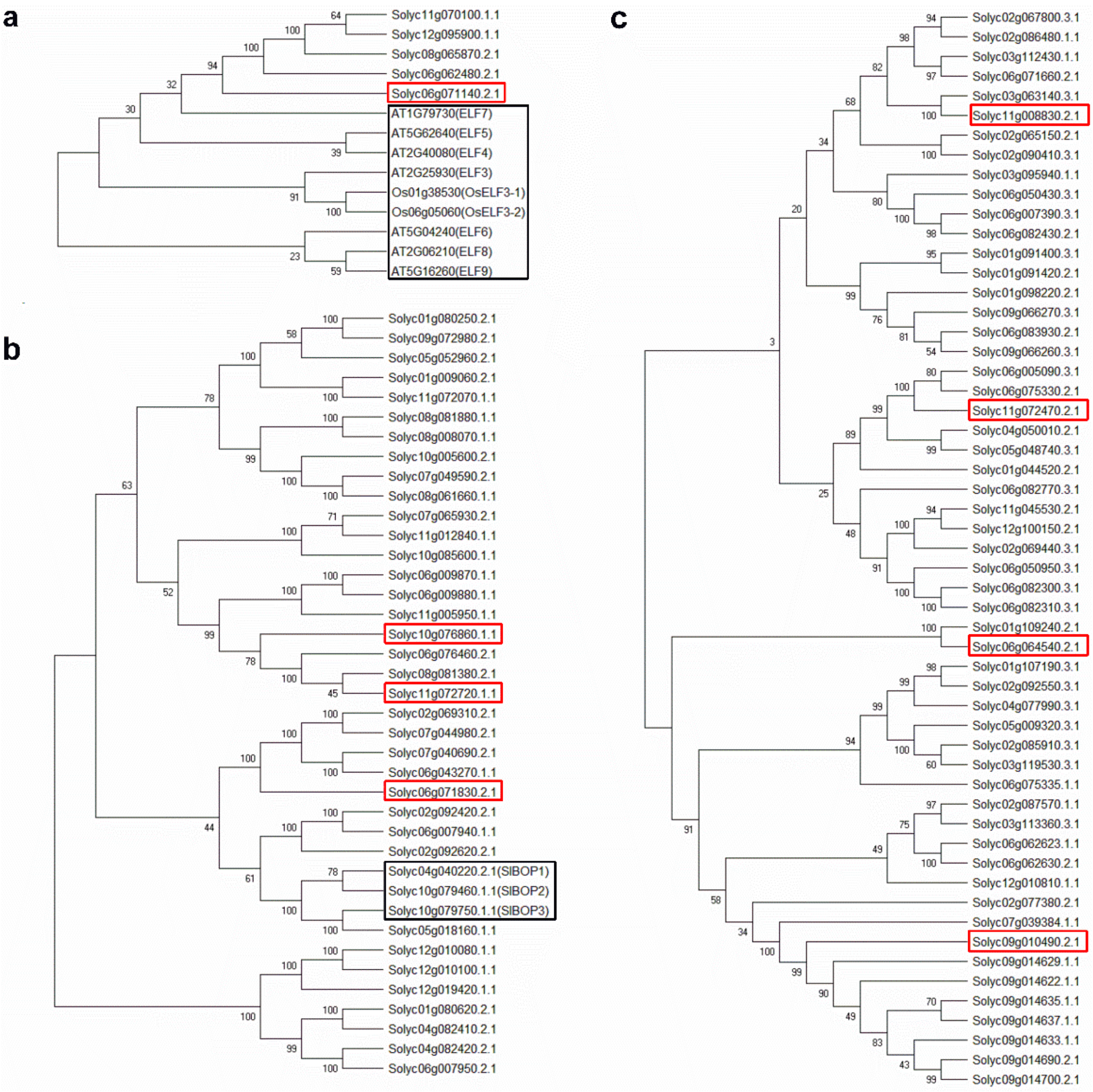


**Supplementary Figure 35: Phylogenetic analysis of ELFs in tomato and *Arabidopsis* (a), BOPs in tomato (b) and LOBs in tomato (c).** Genes in black boxes are those with known functions, and genes in red boxes are the new candidate genes identified by the GWAS. Neighbor-joining trees were constructed using MEGA5. Numbers on the branches indicate bootstrap supports based on 1000 replicates.

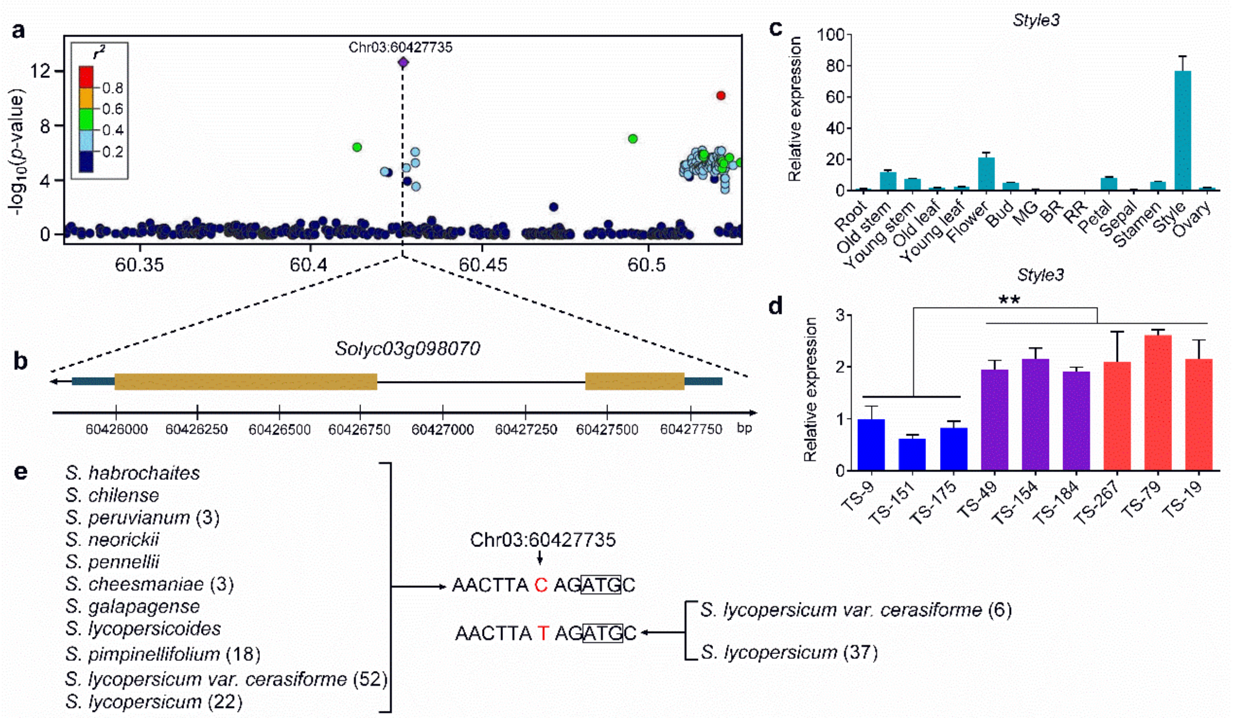


**Supplementary Figure 36: GWAS for stigma exsertion and expression profiles of the candidate gene.** (**a**) Reginal Manhattan plot of GWAS for stigma exertion. The figure shows the genomic region spanning 100 bb on either side of the peak SNP (SL2.50ch03_60427735; indicated in purple) on chromosome 3. Pairwise $r^2$ values (a measure of LD) between the lead SNP and all other SNPs in this 200-kb region are indicated by different colors of the dots. (**b**) Gene structure of *Style3* (*Solyc03g098070*), the closest gene to the lead SNP (2 bp downstream of initiation codon ATG). Filled dark green, filled orange and blacklines represent 3' & 5' UTRs, coding sequence and introns, respectively. (**c**) Transcript levels of *Style3* in different tomato organs of Ts-151 (a stigma inside accession). (**d**) Transcript levels of *Style3* in the style of 9 toamto accessions. Ts-9, Ts-151 and Ts-175 are three stigma inside accessions; Ts-49, Ts-154 and Ts-184 are three stigma flush accessions; and Ts-19, Ts-79 and Ts-267 are three stigma exertion accessions. (**e**) Distribution of the two alleles at the lead SNP locus in different tomatoes. Numbers between brackets indicate the sequences analysed within each species/accession.

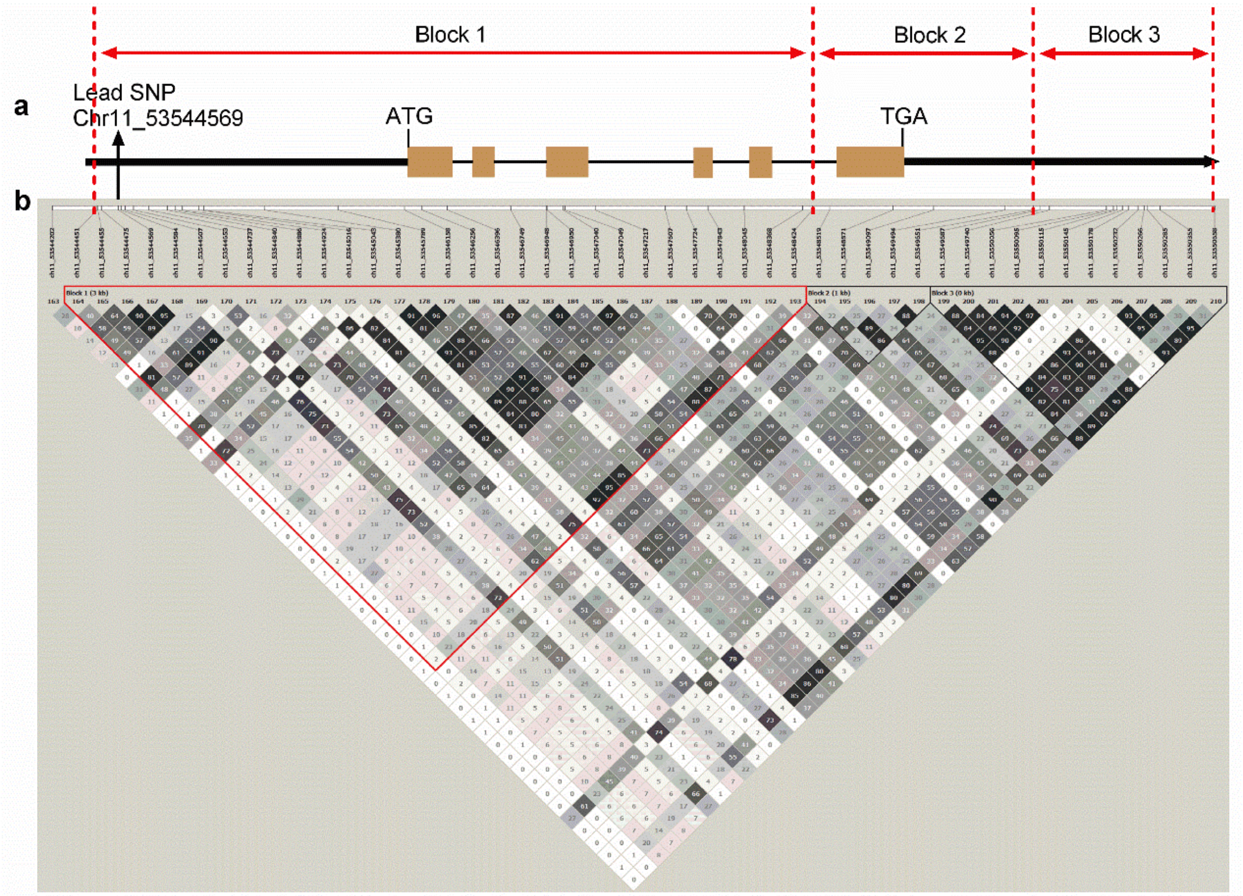


**Supplementary Figure 37: Gene structure and LD blocks surrounding *SlALMT15*.** (**a**) Gene structure of *SlALMT15*. Filled black, filled orange and blacklines represent promoter & UTRs, coding sequence and introns, respectively. The position of the lead SNP (SL2.50ch11_53544569) is pointed by the black arrow. (**b**) Representation of the pairwise $r^2$ values (a measure of LD) among all polymorphic sites in genomic region corresponding to (**a**). The 3 haploblocks are presented in which block 1 containing the lead SNP is indicated by red inverted triangle.

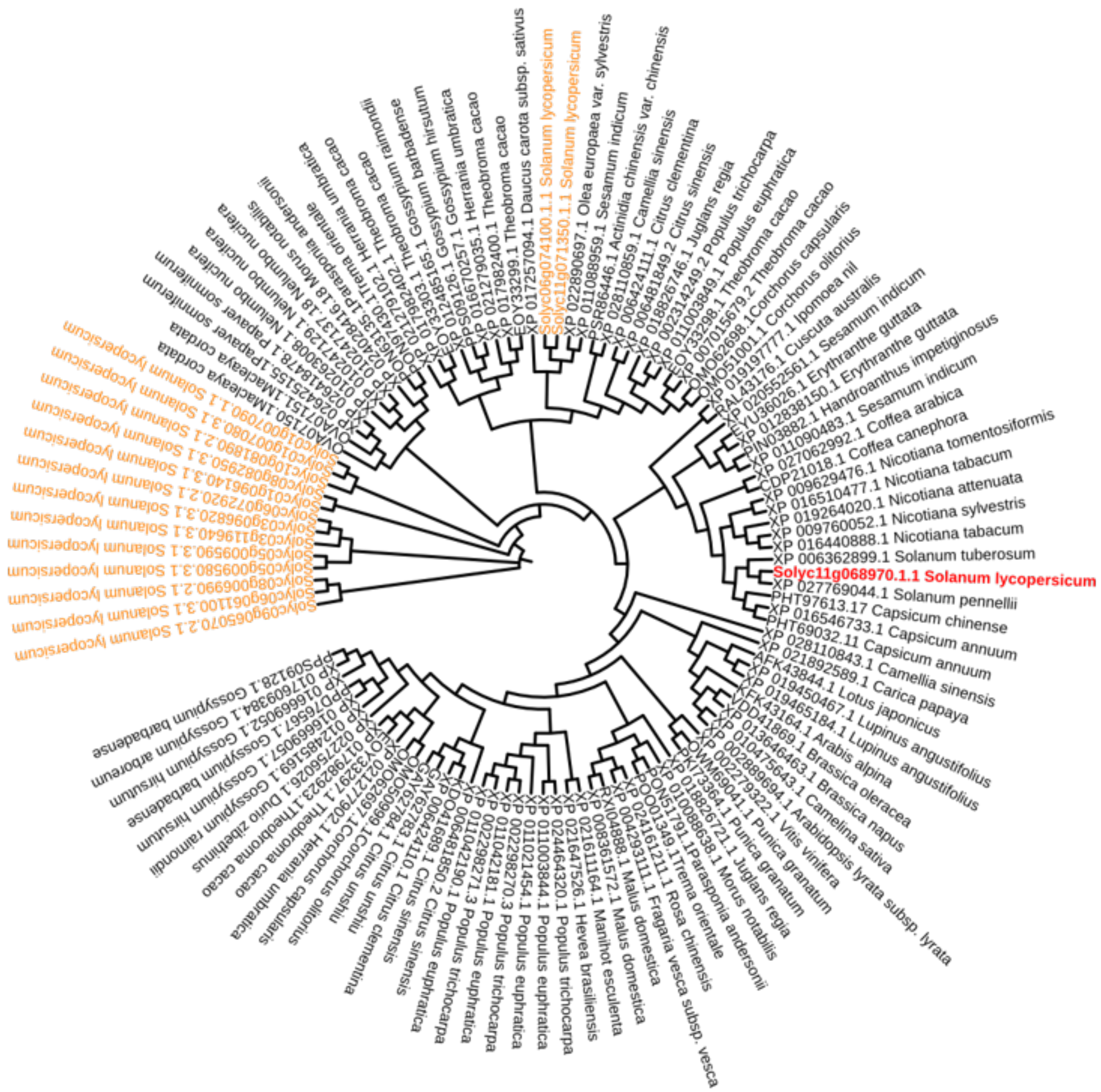


**Supplementary Figure 38: Phylogenetic analyses of SlALMT15.** Phylogenetic analysis of ALMTs in different species. Full-length sequences of *SlALMT15* orthologs from various plants were collected following NCBI-BLAST. The neighbor-joining tree was constructed using MEGA5. SlALMT15 is shown in red, its paralogs in tomato are shown in orange and its orthologs in other plant species are shown in black.

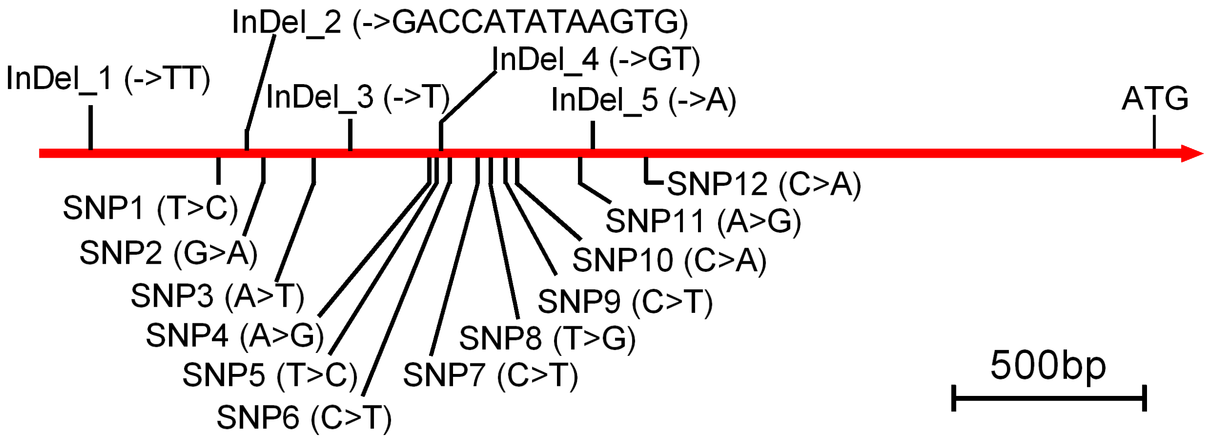


**Supplementary Figure 39: Variations in the *SlALMT15* promoter among 13 tomato accessions with different stomata densities.** The 5 InDels and 12 SNPs are identified and shown. ATG means the initiation codon of *SlALMT15*.

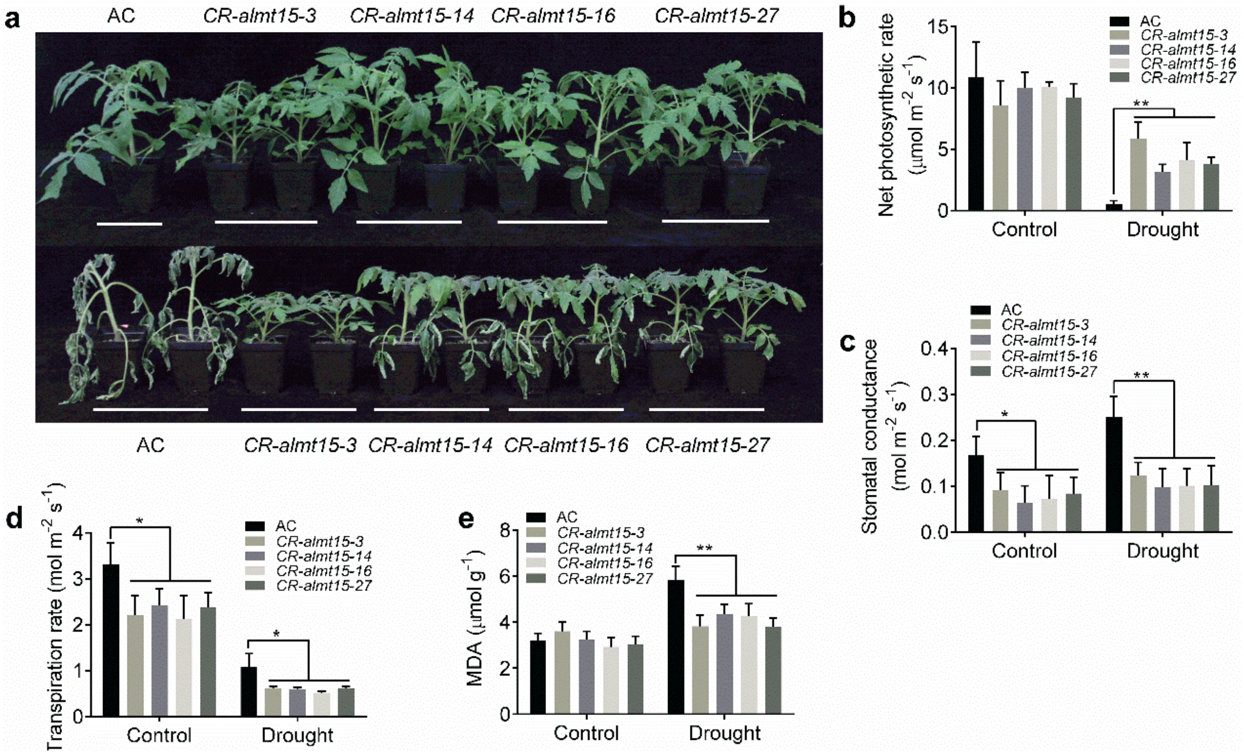


**Supplementary Figure 40: CRISPR/Cas9-engineered mutations in *SlALMT15* result in enhanced drought tolerance in tomato.** (**a**) Phenotypes of seedlings from almt15 mutants and wild-type under drought stress conditions. Five-week old seedlings from $T_2$ *SlALMT15* knockout lines and wild-type were subjected to drought stress by withholding water for 8 d. (**b-e**) Net photosynthetic rates (**b**), transpiration rates (**c**), stomatal conductance (**d**) and MDA levels (**e**) in leaves of *SlALMT15* knockout lines and wild-type plants grown in normal and drought stress conditions (n=3). *, $P < 0.05$; **, $P < 0.01$, Student's *t*-test.

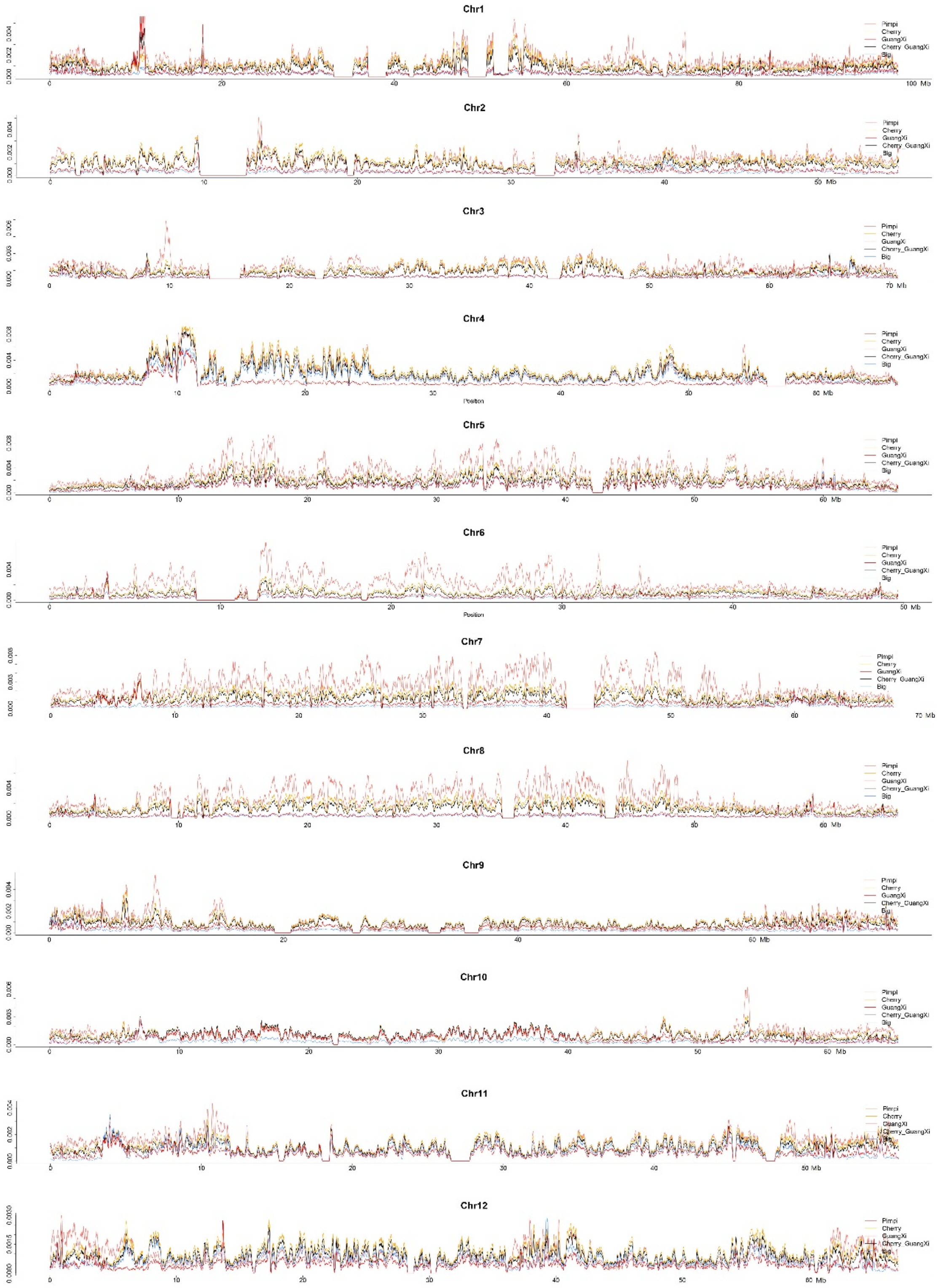


**Supplementary Figure 41:** Distribution of nucleotide diversity (π) for SPIM (light pink), SLC (orange), GX (red), SLC_GX (black) and SLL (light blue) across the 12 chromosomes.
