## Supplementary Note for "Genome-wide association study reveals the genetic architecture of 27 yield-related traits in tomato"

**Methods and description for phenotype evaluation**

1. **FIN (first inflorescence node):** Number of leaves from cotyledon to the first inflorescence on the main stem. Six plants for each accession were investigated in each biological repeat.
2. **FSIN (first to second inflorescence node):** Number of leaves from the first inflorescence to the second inflorescence on the main stem. Six plants for each accession were investigated in each biological repeat.

3. **IL (internode length):** Length (cm) of the main stem between the second inflorescence to the third inflorescence. Six plants for each accession were investigated by vernier caliper in each biological repeat.

4. **SP (self pruning):** Depending on whether leaf or flower buds are differentiated from the stem apical meristem, tomato can be divided into infinite growth type and finite growth type. According to the following figure, data were recorded as ‘0’ or ‘1’.

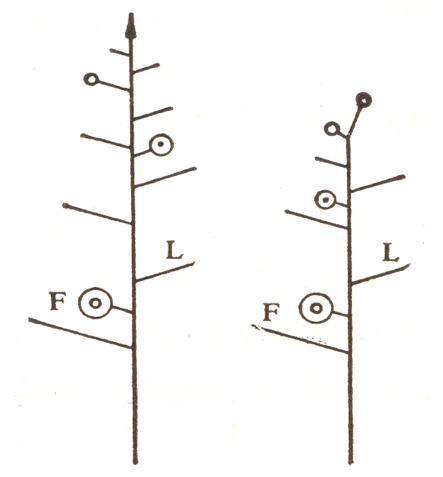

**b**

**a**

Figure m1 Different growth habits of tomatoes

a: ‘0’, infinite growth type; b: ‘1’, finite growth type

5. **FB (flower branch):** According to leaf buds that are differentiated or not on the inflorescence, data were recorded as ‘0’ and ‘1’.

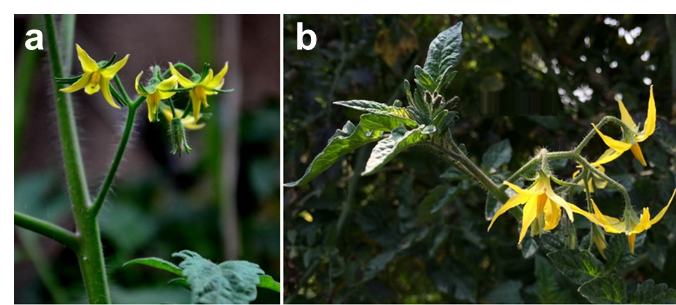

Figure m2 Different type of tomato inflorescence branch

a: ‘0’, none; b: ‘1’, appearance

6. **FLNS (flower number on the second inflorescence):** Number of flowers on the second inflorescence at the full-bloom stage (>90% flower opened in the second inflorescence). Six plants for each accession were investigated in each biological repeat.
7. **FLNT (flower number on the third inflorescence):** Number of flowers on the third inflorescence at the full-bloom stage (>90% flower opened in the third inflorescence). Six plants for each accession were investigated in each biological repeat, and the average number was calculated.
8. **FRNS (fruit number on the second truss):** Number of fruits on the second truss at the mature green stage (the first fruit begin to mature on the second truss). Six plants for each accession were investigated in each biological repeat, and the average number was calculated.
9. **FRNT (fruit number on the third truss):** Number of fruits on the third truss at the mature green stage (the first fruit start to mature on the third truss). Six plants for each accession were investigated in each biological repeat, and the average number was calculated.
10. **IT (inflorescence type):** According to the following pictures, based on the number of inflorescence branches, data were recorded as ‘1’, ‘2’, ‘3’ and ‘4’

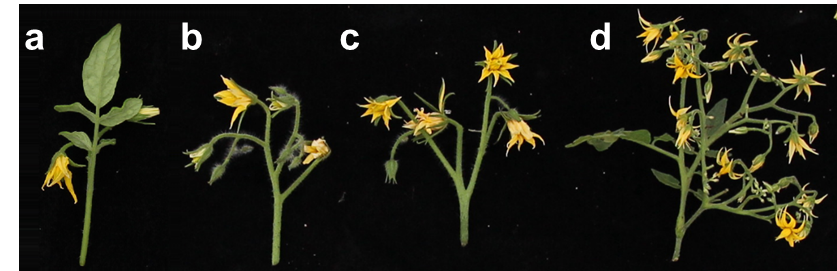

Figure m3 Different types of tomato inflorescence type

a: ‘1’, signal flower type; b: ‘2’, simple inflorescence type; c: ‘3’, double inflorescence type; d: ‘4’, compound inflorescence type

11. **FL (fasciated flower):** >5% flowers were fasciated on the second inflorescence at the full-bloom stage (>90% flower opened on the second inflorescence). According to the following pictures, data were recorded as ‘0’, and ‘1’

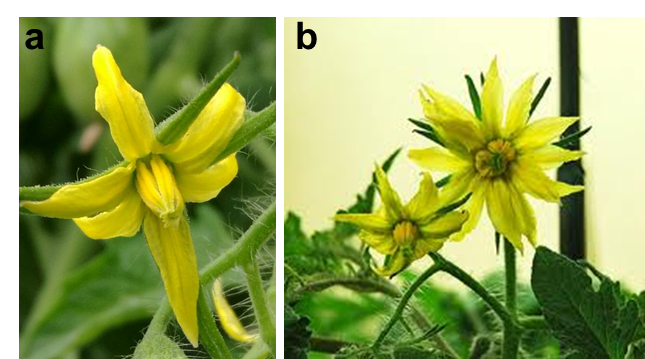

Figure m4 Different type of tomato flower

a: ‘0’, normal flower; b: ‘1’, fasciated flower

12. **PN (****petal number):** Number of petals on each flower (Figure m5). Ten flowers were collected for each accession in each biological repeat, and the average number was calculated.
13. **SN (sepal number):** Number of sepals on each flower. Ten flowers were collected for each accession in each biological repeat, and the average number was calculated.
14. **PL (****petal length):** Length (mm) of the petal (Figure m5). Ten flowers were collected for each accession in each biological repeat, and the average length (measured by vernier caliper) was calculated.

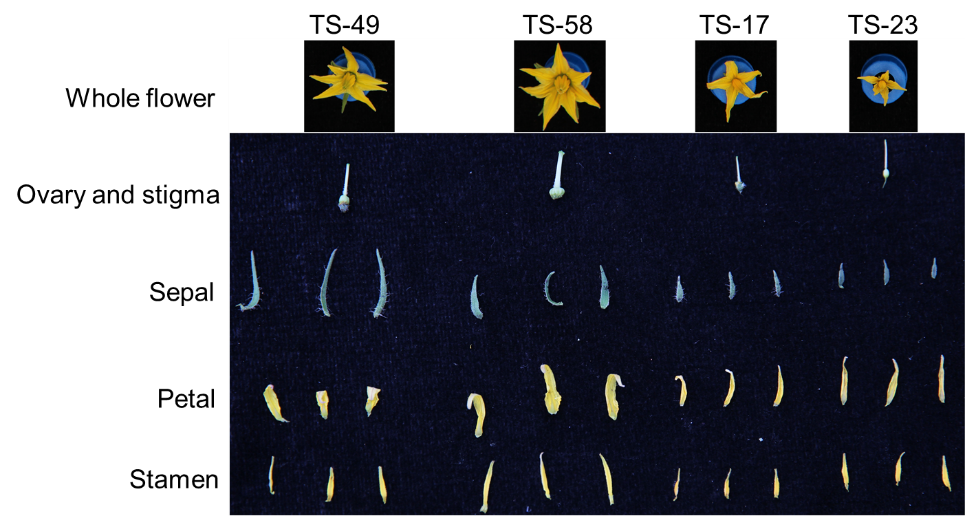

Figure m5 Flower -related floral organ phenotype

15. **SL (sepal length):** Length (mm) of the sepal (Figure m5). Ten flowers were collected for each accession in each biological repeat, and the average length (measured by vernier caliper) was calculated.
16. **SPR (sepal length to petal length ratio):** Ratio of sepal length to petal length.
17. **OLD (ovary longitudinal diameter):** Longitudinal diameter (mm) of ovary. Ten flowers were collected for each accession in each biological repeat, and the average length (measured by vernier caliper) was calculated.
18. **OTD (ovary transverse diameter):** Transverse diameter (mm) of ovary. Ten flowers were collected for each accession in each biological repeat, and the average length (measured by vernier caliper) was calculated.
19. **OTLD (ovary transverse diameter to ovary longitudinal diameter ratio):** Ratio of ovary transverse diameter to ovary longitudinal diameter.
20. **STAL (stamen length):** Length (mm) of the stamen (Figure m5). Ten flowers were collected for each accession in each biological repeat, and the average length (measured by vernier caliper) was calculated.
21. **STIL (stigma length):** Length (mm) of the stigma (Figure m5). Ten flowers were collected for each accession in each biological repeat, and the average length (measured by vernier caliper) was calculated.
22. **SSR (****stamen length to stigma length ratio):** Ratio of stamen length to stigma length.
23. **SE (****stigma exsertion):** According to the following pictures, based on the relative length of stamen and stigma, data were recorded as ‘1’, ‘2’ and ‘3’

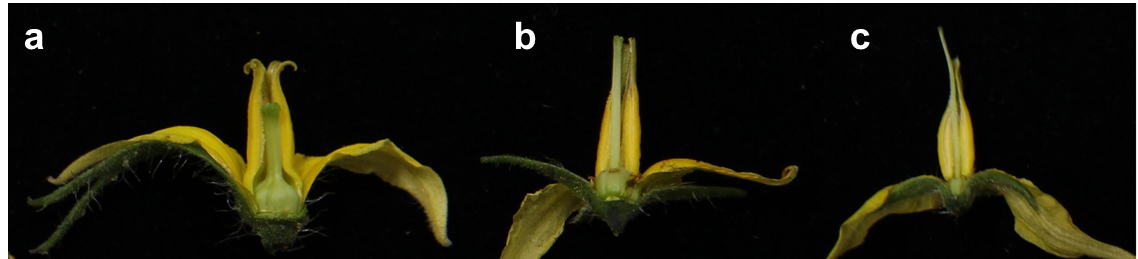

Figure m6 Different types of stigma exsertion

a: ‘1’, stigma shorter than stamens; b: ‘2’, stigma is as long as the stamen; c: ‘3’, stigma longer than stamens

24. **SS (stigma shape):** According to the figure m6, the stigma shape was recorded as ‘1’: single round style stigma (Figure m6c), ‘2’: flat style stigma (Figure m6b) or ‘3’: split style stigma (Figure m6a).
25. **FSD (fruit stalk diameter):** According to the figure m7, the diameter of fruit stalk was measured by vernier caliper (mm). Ten fruits (at the mature green stage) were collected for each accession in each biological repeat, and the average length was calculated.

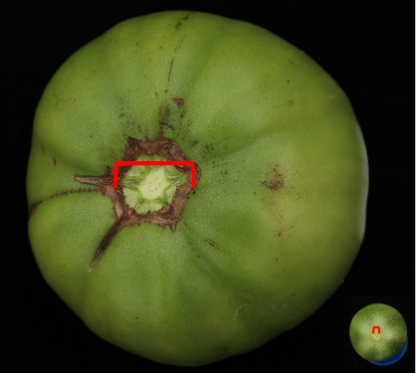

Figure m7 Diversity of fruit stalk. Red mark indicates the diameter of the fruit stalk.

26. **FSL (fruit stalk length):** According to the figure m8, for jointless accessions, FSL was measured from the point of attachment on the stem to the base of the calyx. For joint accessions, FSL was measured from the point of joint to the base of the calyx. The length of fruit stalk was measured by vernier caliper (mm). Ten fruits (at the mature green stage) were collected for each accession in each biological repeat, and the average length was calculated.

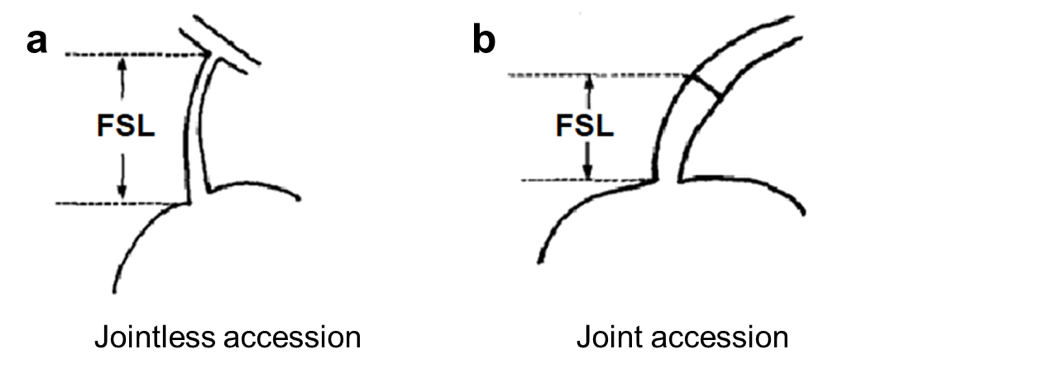

Figure m8 Fruit stalk length for jointless accessions (a) and joint accessions (b)

27. **SD (stomatal density)**: For each accession, the upper third fully expanded leaves from five-week-old seedlings were collected for stomatal density measurement. The stomata were counted in each field of view (200×: ~0.065 mm^2^) using an Olympus BHS/BHT System microscope (BH-2), Three fields per leaf were observed for each accession, and the average number was calculated.

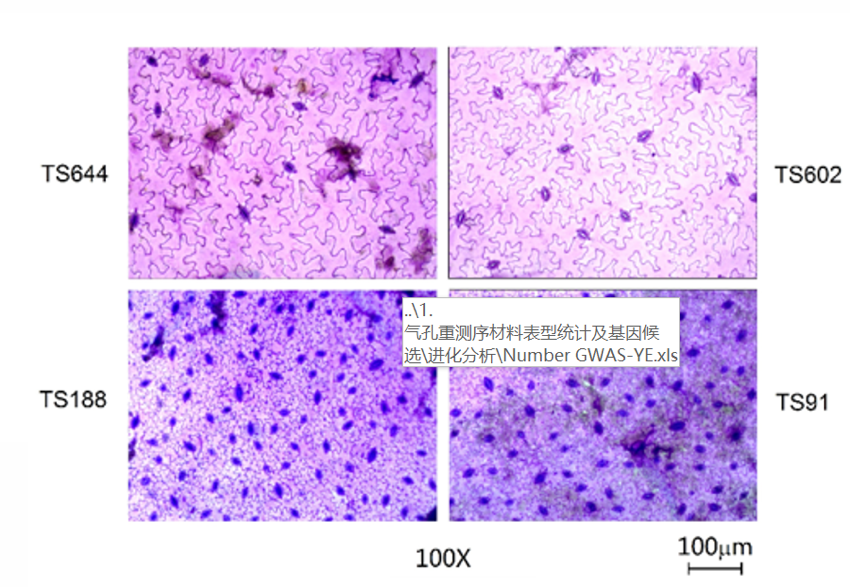

Figure m9 Stomatal density in different tomato accessions
